# Rapid, systematic updating of movement by accumulated decision evidence

**DOI:** 10.1101/2023.11.09.566389

**Authors:** Manuel Molano-Mazón, Alexandre Garcia-Duran, Jordi Pastor-Ciurana, Lluís Hernández-Navarro, Lejla Bektic, Debora Lombardo, Jaime de la Rocha, Alexandre Hyafil

## Abstract

Acting in the natural world requires not only deciding among multiple options but also converting decisions into motor commands. How the dynamics of decision formation influence the fine kinematics of response movement remains, however, poorly understood. Here we investigate how the accumulation of decision evidence shapes the response orienting trajectories in a task where freely-moving rats combine prior expectations and auditory information to select between two possible options. Response trajectories and their motor vigor are initially determined by the prior. Rats movements then incorporate sensory information as early as 60 ms after stimulus onset by accelerating or slowing depending on how much the stimulus supports their initial choice. When the stimulus evidence is in strong contradiction, rats change their mind and reverse their initial trajectory. Human subjects performing an equivalent task display a remarkably similar behavior. We encapsulate these results in a computational model that, by mapping the decision variable onto the movement kinematics at discrete time points, captures subjects’ choices, trajectories and changes of mind. Our results show that motor responses are not ballistic. Instead, they are systematically and rapidly updated, as they smoothly unfold over time, by the parallel dynamics of the underlying decision process.

## Introduction

Grabbing an apple at the grocery store requires not only selecting the best one, but also planning the arm’s trajectory to reach it. In recent years, the classical view of the brain working sequentially, first deciding what to do and then deciding how to do it, has been revised in favor of a perspective that describes these two processes as running in parallel (Gallivan et al. 2018; Wispinski, Gallivan, and Chapman 2020). This new perspective allows not only to predict motor features (the *how to do it*) from cognitive variables (that determine *what to do*), but also to access the cognitive processes underlying the production of a given behavior through the study of motor kinematics (Seideman, Stanford, and Salinas 2018; Korbisch et al. 2022). Motor responses supporting a choice have been characterized by their speed of execution (i.e. vigor), and by the existence of changes in motor plans along their execution (i.e. trajectory updates). Vigor is linked to the subjective value of the chosen option (Shadmehr et al. 2019), increasing with reward, both in reaching (Summerside, Shadmehr, and Ahmed 2018) and saccadic responses (Milstein and Dorris 2007; Xu-Wilson, Zee, and Shadmehr 2009). Vigor is also impacted by urgency in humans (Thura 2020) and non-human primates (Thura et al. 2014). However, the effect on response vigor of cognitive factors such as the evidence towards the decision is poorly understood.

Trajectory updates have been observed when sensory evidence is presented sequentially and can be incorporated into an unfolding trajectory (Song and Nakayama 2009; Stone, Mattingley, and Rangelov 2022). Monkeys (Kiani et al. 2014; Kaufman et al. 2015) and humans (Resulaj et al. 2009; van den Berg et al. 2016) often reverse an initial choice if it is contradicted by novel sensory evidence (a *change of mind,* or *CoM*). Yet, we do not know if this online updating of trajectories represents an isolated phenomenon that only occurs when new evidence promotes a categorical change in the initial choice. Alternatively, *all* trajectories could be systematically updated based on novel relevant information, even when the endpoint is not affected. Where the brain operates between these two opposite scenarios has yet to be unveiled.

Our understanding of the internal processes that control decision-making has greatly benefited from computational models such as the Drift-Diffusion Model (DDM) (Roger Ratcliff and McKoon 2008). The DDM summarizes the evidence supporting the different options into a *decision variable* that evolves in time until it reaches a bound that determines the choice. This intuitive paradigm generates precise predictions about how choices and reaction times depend on stimulus evidence (R. Ratcliff 1985; Palmer, Huk, and Shadlen 2005; Bogacz et al. 2010; Pardo-Vazquez et al. 2019) and prior expectations (Bogacz et al. 2006; Roger Ratcliff and McKoon 2008; Urai et al. 2019). Recent work has extended the DDM to account for proactive responses, whereby rats and humans initiate their response at a time that is independent of the stimulus (Hernández-Navarro et al. 2021; Hawkins and Heathcote 2021). This extended model implies that, during these proactive responses, subjects incorporate the stimulus information into their response trajectories as they are moving. However, the DDM and its extensions do not characterize either the response trajectories nor the nature, extent and timing of any interaction between decision variables and trajectory kinematics. Bridging the gap between the internal decision dynamics and the precise movement kinematics requires the development of new models that, capitalizing on the explanatory power of accumulation models, can ultimately describe time-resolved response trajectories.

Here we investigate in rats and humans how the dynamics of evidence accumulation impacts the response trajectories in a prior-guided auditory discrimination task. We track the subjects’ trajectories and find that response vigor is modulated by the amount of accumulated decision evidence provided both by the prior and the stimulus. Subjects incorporate the sensory evidence into their unfolding response trajectory, by adjusting its vigor or by reverting the planned choice when sensory evidence strongly contradicts the initial decision, i.e. performing a change of mind. We encapsulate these observations in a new computational model that simulates both the dynamics of evidence accumulation and the execution of the response. When fitted to subject behavior, the model provides a parsimonious description of the relationship between these two processes, in particular the impact of the accumulated evidence on the response vigor and the conditions yielding changes-of-mind. Altogether, our results show that the evidence accumulated during perceptual decisions controls the kinematics of the response trajectory, systematically updating them *en route* as new evidence arrives.

## Results

### Movement time depends on decision variables

We investigated how response trajectories are formed as rats (Groups 1 and 2; n=15; see Methods) integrate decision evidence in a perceptual task where decisions are guided both by an acoustic stimulus and the recent trial history (Hermoso-Mendizabal et al. 2020) (Figure 1a). On each trial, following a 300-ms fixation period, a stimulus was played from two lateral speakers until the animal poked out from the center port and headed towards one of two side ports. Rats were rewarded if they selected the port associated with the louder speaker (Pardo-Vazquez et al. 2019). The stimulus strength was manipulated by varying the intensity difference between the two speakers. Moreover, trials were organized into repeating and alternating contexts, in which the probability that the stimulus category (i.e. Left vs Right) is the same as in the previous trial was 0.8 and 0.2, respectively (Figure 1a). Rats leveraged these serial correlations in the trial sequence by building a prior expectation of the next rewarded port based on the recent history of responses and outcomes (Hermoso-Mendizabal et al. 2020). On each trial, we estimated the magnitude and category of this subjective prior, a variable we called *prior evidence*, by fitting a logistic regression model (see Methods). Rats’ final choices combined the prior evidence, accumulated across several trials, and the stimulus evidence (Figure 1b).

**Figure 1.**
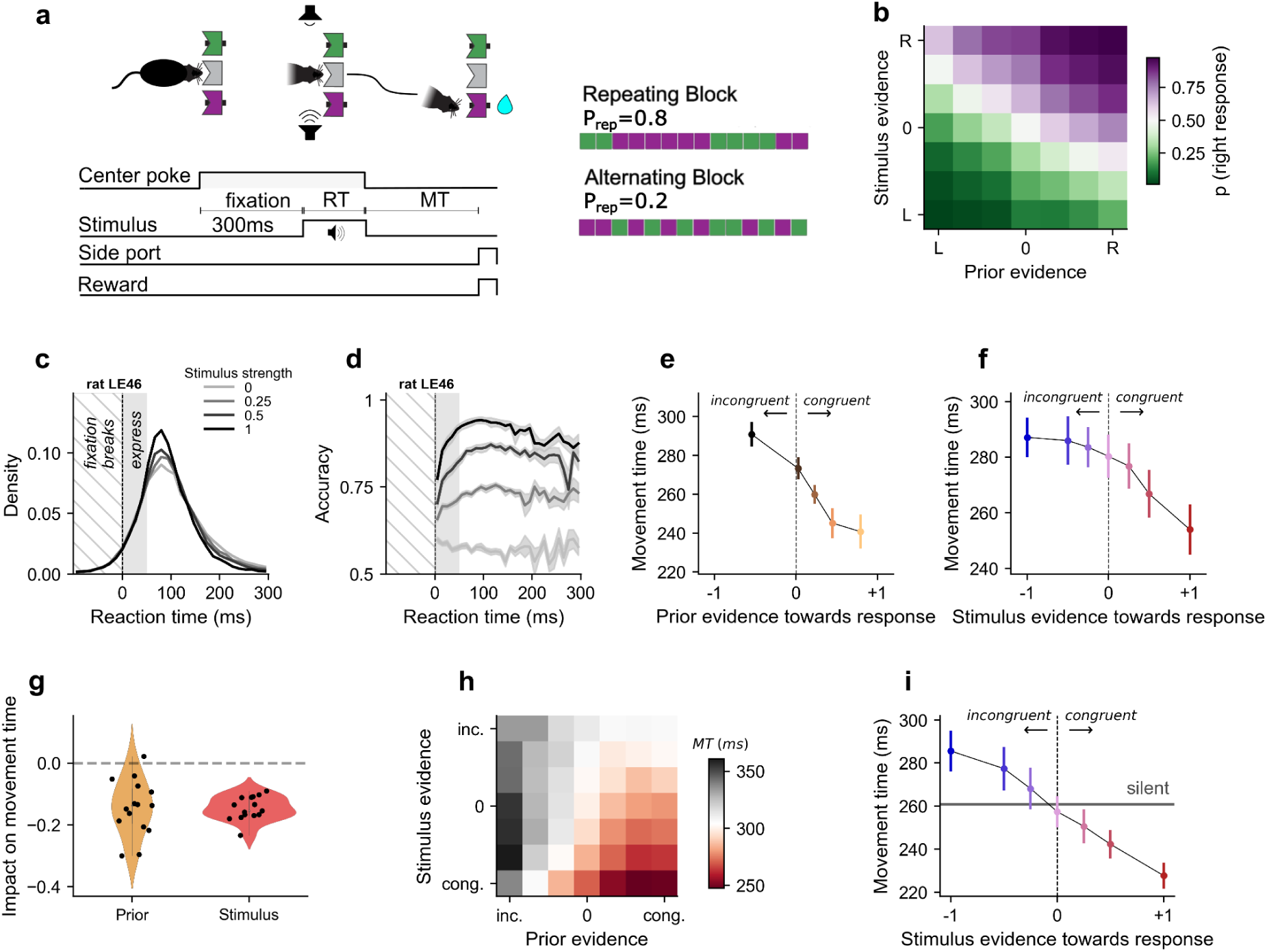
Rats reaction times, choices and motor times in an auditory, prior-guided, two-alternative categorization task. **a)** On each trial, rats fixated for 300 ms in the center port before an acoustic stimulus was played from two lateral speakers. Rats were free to initiate a response toward a lateral port at any time following stimulus onset. RT: reaction time defined from stimulus onset to initiation of rat response (central port withdrawal); MT: movement time defined from response initiation to the lateral port entering). **Right**:Trials are organized into blocks of 80 trials, with the probability of repeating the trial category set to P_rep_=0.8 in repeating blocks and P_rep_=0.2 in alternating blocks. **b)** Proportion of rightward responses as a function of stimulus evidence (i.e. signed stimulus strength) and prior evidence, averaged across animals (Groups 1+2, n=15). **c)** Distribution of reaction times for each stimulus strength, for one typical animal (LE46). Negative RTs correspond to fixation breaks. Shaded area represents express responses (RT<50 ms). **d)** Tachometric curves representing the mean accuracy as a function of RT for the different stimulus strengths, for the same animal. **e)** Median movement time averaged across animals in silent catch trials as a function of prior evidence towards the selected port (i.e. positive if the prior points towards the animal decision and negative otherwise; binned in 5 quantiles). Error bars: standard error of the mean (s.e.m.). **f)** Median movement time in express response trials (RT< 50 ms) with small prior (|z| < 10% percentile), as a function of stimulus evidence towards the selected port. **g)** Regression weights of the movement time against the prior and stimulus evidence towards the response. Each dot represents one animal. **h)** Mean movement time as a function of prior and stimulus evidence towards the selected port, averaged across animals. **i)** Average movement time as a function of stimulus evidence supporting the decision, in express response trials (RT<50 ms) with prior evidence supporting the decision (prior evidence towards response z > median). Stimuli can both accelerate or slow down the default trajectory revealed in silent trials with the same prior (horizontal line).

Importantly, in our experimental paradigm, the initiation of rat responses was often driven by a proactive process, leading to a large fraction of reaction times (RTs) that were too short to be triggered by the stimulus. *Express responses* (RT< 50 ms) were observed in 34.3 ± 12.2% of all trials (mean±SD; Figure 1c) (Hernández-Navarro et al. 2021). Although the response in these trials was initiated independently from the stimulus, the final choice made by the rat integrated the stimulus evidence, as demonstrated by the increase in accuracy with stimulus strength (Figure 1d). This implies that in a large fraction of trials, rats process the stimulus and update their decision while moving from the central port to the selected lateral port. We therefore set out to characterize how the prior and simulus are dynamically integrated into the unfolding trajectory.

We first asked whether the prior and stimulus evidence impacted the response vigor, that is the velocity with which rats executed their orienting movements to the side port. For each trial, we computed the Movement Time (MT), defined as the time between the poke out from the center port and the poke in a lateral port (Figure 1a). To isolate the contribution of the prior evidence on vigor, we introduced *silent catch trials,* where no acoustic stimulus was played (Group 2, n = 6, mean±SD 6.7 ± 0.6% of all trials). Prior evidence made the responses faster when it pointed towards the selected port: a strong prior congruent with the response shortened MT (congruent prior; Figure 1e). Inversely, when the prior evidence was incongruent with the response, the animal took longer (incongruent prior; Figure 1e). We then analyzed the impact of stimulus evidence on MT, restricting the analysis to trials with small prior magnitude to isolate stimulus influence (see Methods). Similarly to the impact of prior evidence, rat movement responses were faster when the stimulus supported the selected port (congruent stimulus), and slower when the stimulus contradicted it (incongruent stimulus; Figure 1f). When assessed altogether, MT depended on a linear combination of stimulus and prior evidence towards the response (Figure 1g-h). This influence was present in addition to a within-session slowing due to satiation and tiredness (Supplementary Fig. 1) (Hernández-Navarro et al. 2021). These results highlight the relative impact of congruent versus incongruent stimuli. However, they do not clarify whether the net effect of the stimulus on MT was positive or negative, i.e. whether the stimulus accelerates or slows down the initial trajectory. For this, we assessed the impact of sound on MT with respect to silent trials: MT decreased for congruent stimuli, but increased when the stimulus was at odds with the final choice (Figure 1i). This shows that, on average, incongruent stimulus evidence effectively slowed the response originally planned, while congruent evidence sped it up. In summary, the time taken by the rats to perform a response trajectory in a decision-making task displays clear signatures of the decision variables.

### Prior and stimulus evidence impact rats’ trajectory at different times

The above results show that both prior and stimulus evidence impact rats’ response trajectories. To finely characterize this influence, we next studied the impact of these factors on time-resolved response trajectories. We extracted the response trajectories described by the rats during the task from video recordings of a camera placed above the animal. Using an automatic pose estimation method (Mathis et al. 2018), we inferred the coordinates of the snout of the rat in the horizontal plane, for each video frame (Figure 2a; see Methods). We focused our analyses on the dimension aligned with the three ports (Figure 2b), since it contains the most relevant information about the response orienting trajectory. Trajectories along this axis revealed relatively stereotyped movements consisting in smooth sigmoidal-like curves starting at the central port and ending at one of the side ports (Figure 2c). Consistent with our analyses of MT, prior and stimulus evidence accelerated or decelerated the trajectories depending on their congruence with the final choice while preserving the same stereotyped shape (Figure 2d-e). This caused the peak velocity of the animal’s movement to show a similar dependence on the prior and stimulus evidence as the MT (Figure 2f-g).

**Figure 2.**
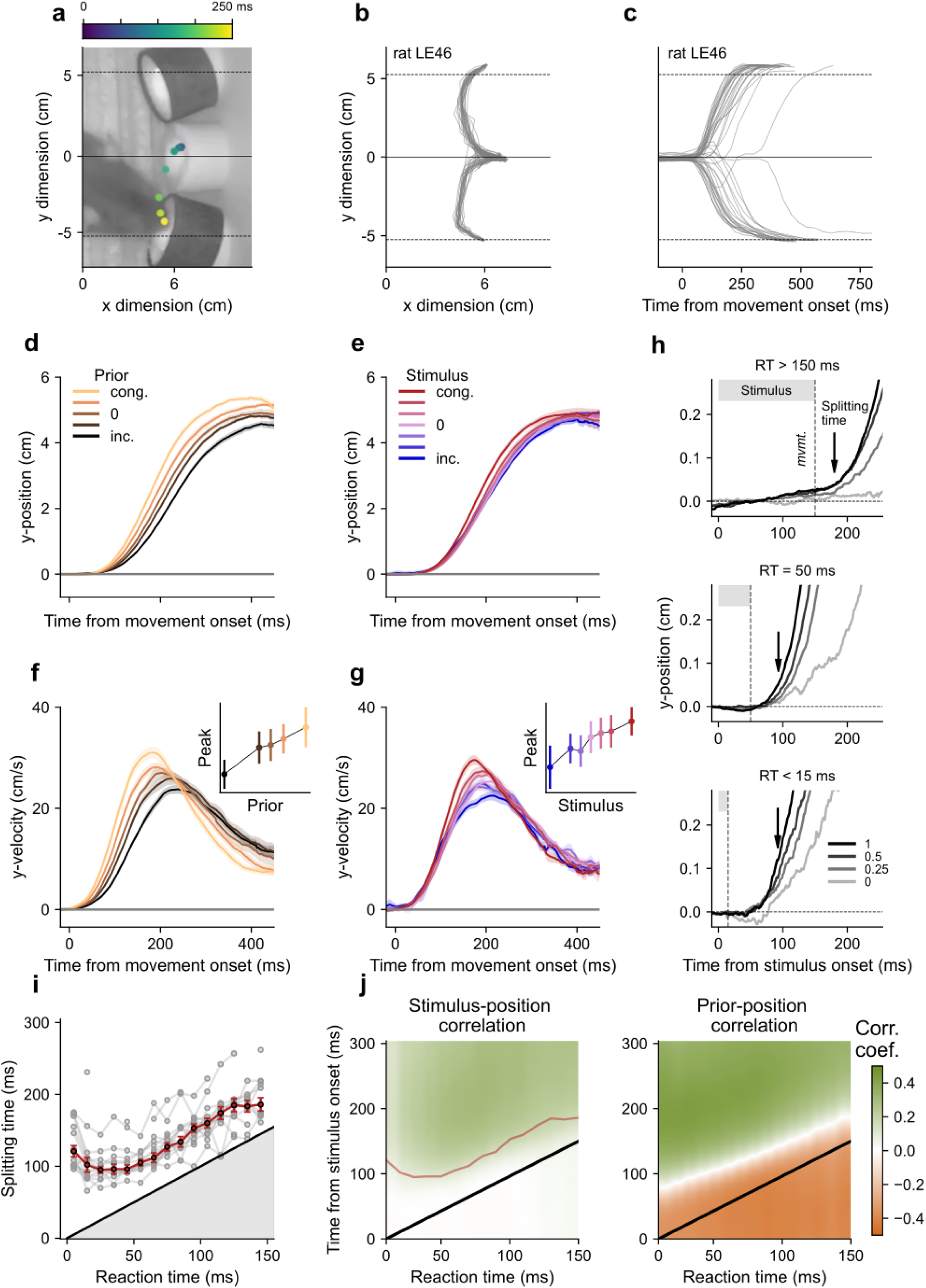
Prior and stimulus evidence modulate response trajectories, at different times. **a)** Example video frame showing a rat entering the right lateral port. Color dots indicate the position of the rat snout, obtained from automatic video analysis, for 6 frames prior to the frame shown (colors correspond to the time elapsed since movement onset, see colorbar). **b)** Trajectories from individual trials. Dashed lines as in panel a**. c)** Position of the rat along the y-dimension as a function of time from movement onset. Dashed lines as in panels a, b. **d, f)** Average position (d) and velocity (f) in silent trials as a function of time from movement onset and prior evidence towards the selected port. Inset in f: average peak velocity versus prior. Trajectories corresponding to rightward responses were flipped along the y axis so that all trajectories end at a positive value. **e, g)** Average position (e) and velocity (g) conditioned on stimulus evidence in express response trials with small prior. Inset in g: average peak velocity versus stimulus evidence. **h)** Splitting time calculation in one example animal. Average trajectories for each stimulus strength in different RT windows: very short (RT<15 ms, bottom), medium (RT between 45 and 55 ms, middle) or long (RT>150 ms, top) reaction times. **i)** Splitting time for stimulus evidence as a function of RT (bins of 10 ms; red trace: median over rats, n=15; gray traces: individual rats; error bars: s.e.m.). Shaded area corresponds to times before the rats poked out of the central port. **j)** Correlation coefficient between the position of the animal and either the stimulus evidence (left panel) or the prior evidence (right panel), as a function of the reaction time (x-axis) and time from stimulus onset (y-axis). Correlation coefficients were averaged across animals. The prior evidence and animal position are negatively correlated before the animal leaves the port, and become positively correlated right after leaving the port. The red curve in the left panel represents the average splitting time (see panel i). The black line represents the RT. Panels d, f show averages across rats for Group 2 (n=6) and panels e, g-j, for Groups 1 (n=9) and 2 (n=6). Error bars in d-g (bands) represent s.e.m..

The impact of the prior onto trajectories occurred earlier than the impact of the stimulus (Figure 2d-g): while the prior information is available at the trial start, sensory evidence must be integrated as the stimulus is played. The precise timing of the stimulus impact hinges on the duration of a sensorimotor processing pipeline that includes the afferent (sensory) delay, evidence integration and efferent (motor) delays. To finely quantify the onset of this impact, we identified the earliest time point from the stimulus onset at which the stimulus strength started affecting the trajectory (Figure 2h; see Methods). This Splitting Time naturally depends on the reaction time, which caps how much processing can be done along the sensorimotor pathway before the animal pokes out from the central port. At long RTs, animals have time to accumulate enough stimulus evidence, form a decision and command the corresponding movement. Therefore, the trajectories generated after long RTs split immediately after poking out, and thus the splitting time approximates the reaction time (after a fixed minimal delay required to detect a significant difference in the trajectories) (Figure 2i). In contrast, at short RTs, the splitting time reaches a plateau: no matter how early the animal leaves the port, it needs a minimal amount of time before incorporating stimulus evidence into the trajectory. This minimum splitting time occurred less than 100 ms after stimulus onset (96.6 ± 25.1 ms across animals), and could be as low as 60 ms in some rats. At very small RTs (RT<10 ms), the stimulus was so short that its impact on the trajectory vanished, causing larger splitting times. The prior information, on the other hand, modulated the position of the animal even before leaving the central port (Figure 2j, right panel). Interestingly, rats prepared their response while inside the central port, by placing their snouts opposite to the side they planned to choose. In summary, the orienting trajectories of the rats were influenced by prior expectations before leaving the port and by stimulus evidence as early as 60 ms after stimulus onset, reflecting the difference in timing of these two sources of information.

### Incongruence between the prior and stimulus evidence leads to changes of mind

These results imply that trajectories, far from being predefined ballistic movements, are updated during their execution based on new sensory evidence. Previous studies in humans and non-human primates have shown that fluctuations in the stream of sensory evidence can lead to changes of mind (CoMs), i.e. a change in the response trajectory towards the option that is aligned with the most recent evidence (Resulaj et al. 2009; Kiani et al. 2014; van den Berg et al. 2016; Peixoto et al. 2021; Boyd-Meredith et al. 2022). We thus wondered whether, in our task, stimuli strongly contradicting the prior-guided initial trajectory could lead rats to perform CoMs, instead of simply slowing down the response without affecting the target port. To identify possible CoMs we selected trials where the trajectory crossed a fixed threshold towards one of the lateral ports but then reversed and ultimately reached the opposite port (Figure 3a-b). This procedure detected 31277 trajectory reversals across all rats (Groups 1 and 2, n = 15), which represent a low but systematic percentage of all trials (2.37 ± 0.95 %, mean±SD across animals; Figure 3c). Although this percentage was sensitive to the value of the threshold used to identify the reversals, the following characterization was not. Trajectory reversals were associated with a stereotyped movement showing a standard initial trajectory towards one side followed by a clear deflection towards the final choice (Figure 3e-f and Supplementary Video 1). The reversal point of the trajectory took place on average 189 ± 15 ms after movement onset and was rarely further than 10 pixels away (∼ 0.7 cm) from the central port (i.e. less than 15% of the distance between central and lateral ports), meaning that rats reversed their trajectories when they had not departed too far from the central port (Figure 3d). Movements associated with a trajectory reversal were on average 129 ms slower than non-reversal responses (Figure 3g), reflecting the time cost associated with longer pathways and the change of direction inherent to those trials.

**Figure 3.**
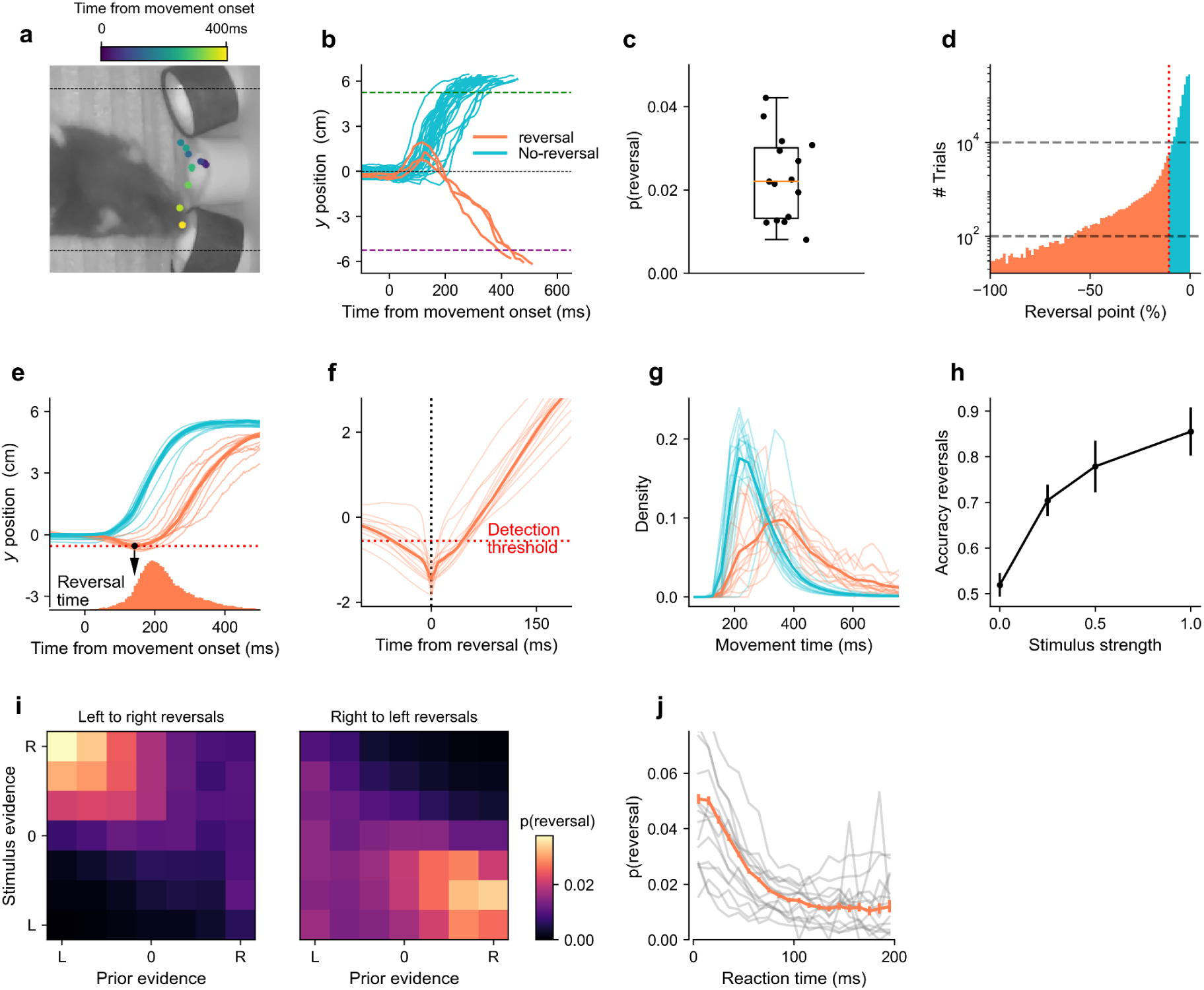
Rats reverse incorrect, prior-driven, initial responses based on incoming sensory information. **a)** Example video frame in a reversal trial: the rat initiates a response to the left port but then reverses and heads towards the right port. Legend as in Figure 2a. **b)** Reversal trajectories (orange traces) and standard trajectories (i.e. no reversal, blue traces) showing the position of the animal snout along the y-axis across time, from an example session. Response ports are represented with dashed lines. Only the left-response no-reversals and right-response reversals in 100 randomly chosen trials from an example session are shown. **c)** Proportion of reversals for individual rats (Groups 1-2, n=15). **d)** Histogram of the reversal point position, i.e. the maximum distance from the central port towards the non-selected port (expressed as a percentage of the distance from the central to the lateral ports), obtained from all animals. Trials with reversal points larger than the detection threshold (vertical red dashed line) are marked as trajectory reversals. **e)** Average trajectories for reversal and standard trajectories aligned at movement onset for individual animals (thin lines) and averaged across animals (thick lines). Bottom: Distribution of the reversal time defined as the time from motion onset to the reversal point. **f)** Average trajectories for reversals aligned to the reversal time. Red dashed line indicates the reversal threshold used to detect reversals (8 pixels, i.e. 0.56 cm). **g)** Distribution of movement time for reversal (orange) and no-reversal trajectories (blue), for individual animals (thin) and averaged across animals (thick). Mean ± SD movement time for non-CoM trials was 283 ± 31 ms and for CoM trials was 421 ± 61 ms. **h)** Mean choice accuracy of reversal trials as a function of stimulus strength, averaged across animals. **i)** Proportion of left-to-right reversals and right-to-left reversals as a function of stimulus and prior evidence (binned in quantiles), averaged across animals. **j)** Proportion of reversals as a function of reaction time (gray traces: individual rats; orange trace: mean ± s.e.m.).

Several characteristics of the trajectory reversals suggest that they corresponded to changes of mind, whereby a first decision driven by the early evidence (provided by the prior) is later switched based on novel (sensory) evidence contradicting it. First, trajectory reversals were usually corrective, i.e. they amended a mistake incurred by the animals’ initial decision, and they were more likely to be so for stronger stimuli (Figure 3h) (Resulaj et al. 2009). Second, trajectory reversals emerged usually when the prior and stimulus were contradictory, with the initial response aligned with the prior while the final response aligned with the stimulus (Figure 3i), consistently across rats (Supplementary Fig. 2). Finally, reversals were more frequent for short RTs (Figure 3j), when the initial trajectory started too soon to be influenced by sensory information and thus was based only on prior information.

In summary, rats’ orienting responses are routinely updated based on incoming sensory information, which either modulates the response vigor (Figure 2) or promotes a CoM that causes a complete reversal of the initially selected choice (Figure 3).

### Human response movements display similar modulation of movement by decision variables

We next wondered whether these results generalize across species. We instructed a group of human subjects (*n*=14) to perform an intensity discrimination auditory task that closely mimicked the rat paradigm (see Methods). Briefly, subjects reported their response by sliding their finger on a tablet from a fixation point to one of two possible targets (Figure 4a-b). As in the rat’s paradigm, the probability of repeating the previous trial category varied between Repeating and Alternating blocks (changes of blocks were not cued). We explicitly inserted an urgency component into the task, forcing subjects to initiate their response within 300 ms after stimulus onset.

**Figure 4.**
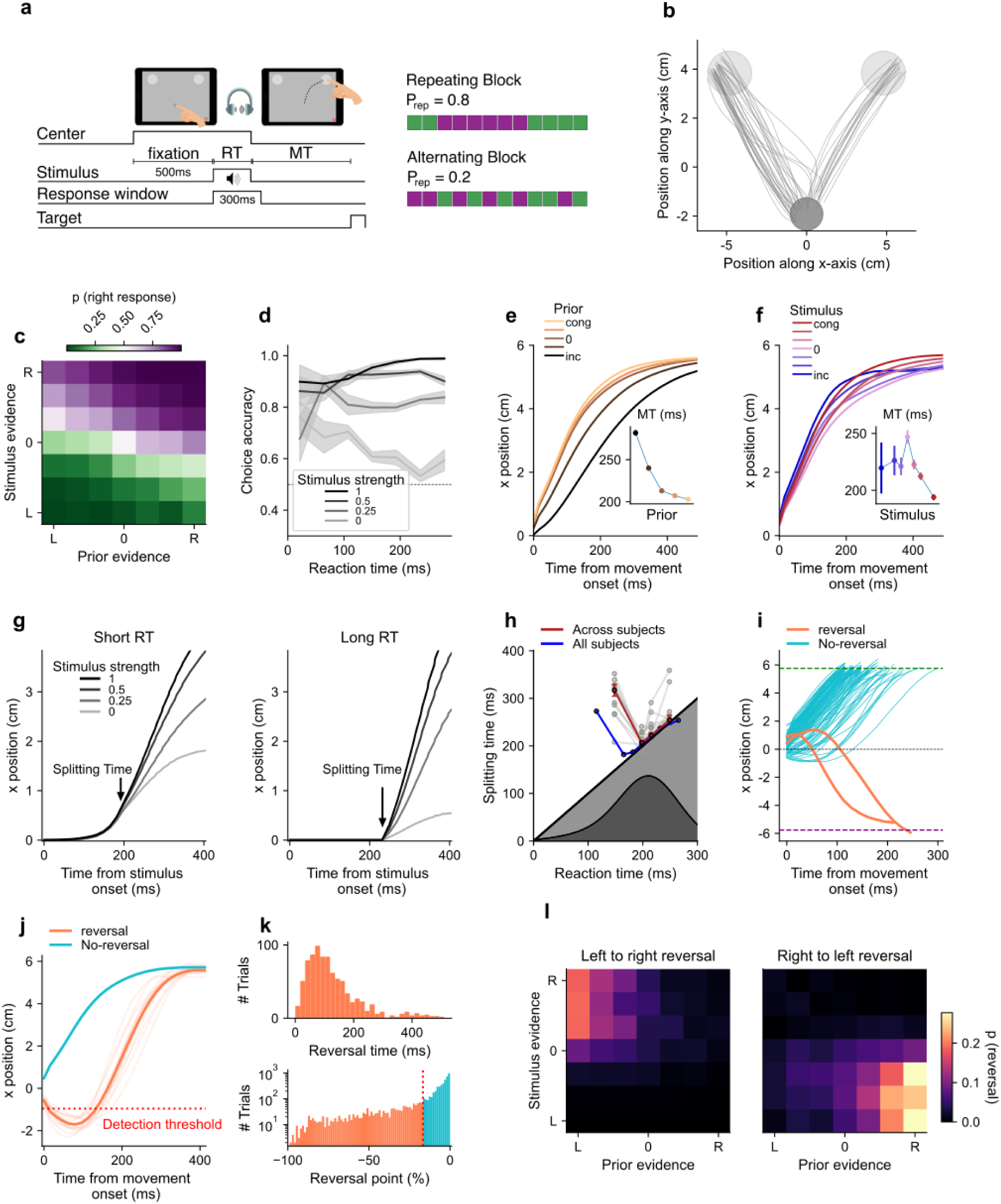
Human response trajectories exhibit similar properties to rats’ trajectories. **a)** Behavioral paradigm. In each trial, human participants (n=14) maintained their index finger fixed on the fixation center circle of a tablet during 500 ms. An acoustic sound was then played through headphones. The participants had to initiate a trajectory along the tablet no later than 300 ms after stimulus onset and reach for one of the two target circles based on the interaural level difference of the stimulus. **b)** Example trajectories from a typical session. **c)** Proportion of rightward responses as a function of stimulus evidence and prior evidence, averaged across subjects (as in Figure 1b). **d)** Mean accuracy as a function of RT for the different stimulus strengths (as in Figure 1d). RT bin width was 43 ms. **e)** Average trajectory as a function of prior evidence towards the selected port. Inset: Average movement time as a function of prior evidence, as in Figure 1e. Legend as in Figure 2d. **f)** Average trajectory as a function of stimulus evidence, in low prior trials. Inset: Average movement time across trials as a function of stimulus evidence towards the selected port, as in Figure 1f. Legend as in Figure 2e. **g)** Average trajectories at each stimulus strength, in short RT trials (first tercile, left) and long RT trials (third tercile, right), grouped over participants. **h)** Splitting time computed for each quartile of RT, for each subject individually (gray), and averaged across subjects (red trace; error bar: s.e.m.). The blue trace shows the splitting time, grouping trials over all subjects (7 RT quantiles). Dark gray: distribution of RT across all trials. **i)** Example trajectories for one participant, showing standard (blue) and reversal (orange) trajectories. Legend as in Figure 3b. **j)** Average trajectory for standard trials and trajectory reversal trials. **k)** Distribution of reversal point position and reversal time across participants. **l)** Proportion of trajectory reversals as a function of prior and stimulus evidence, averaged across participants.

Human behavior in the task was remarkably similar to that of rats. First, subjects’ decisions combined the prior and sensory information (Figure 4c, Supplementary Fig. 3a), and their accuracy increased with stimulus strength even for responses initiated less than 50 ms after stimulus onset (Figure 4d). Second, the MT depended on both prior evidence and stimulus evidence: response trajectories were faster or slower depending on the congruence of these variables with the final response (Figure 4e-f). Third, the impact of stimulus evidence on the trajectory was statistically visible as early as 180 ms after stimulus onset (Figure 4g-h). As in rats, the splitting time was actually slightly smaller for moderate reaction times (around 200 ms) compared to very short reaction times. Fourth, we could detect a reliable proportion of trajectory reversals (7.25 ± 3.55 %, mean±SD across subjects), which were revealed by a deflection in the response trajectory occurring around 156 ± 71 ms after movement onset (Figure 4 i-k). Similar to rats, reversals corrected the decision *en route* mostly when the prior provided strong evidence towards one side but the stimulus indicated the opposite side (Figure 4l); and were often associated with short RTs (Supplementary Fig. 3b). Overall, rats and humans show a very similar relationship between the variables influencing a decision and its actual execution.

### A joint model of decision-making and motor trajectories

We formalized our observations about the impact of the prior and stimulus evidence onto response trajectories into a computational model that ties mechanistically the dynamics of a latent decision variable with the animal’s orienting trajectory (Figure 5). The model sets apart from traditional decision-making models by fully describing the dynamics of the orienting response, and not just its end point. It consists of a decision-making module and a motor module. The decision-making module extends the classical Drift-Diffusion Model whereby a decision variable *x*(t), initialized at a value proportional to the prior evidence, integrates sensory evidence over time until one of two decision bounds is reached and the associated side is selected (Good 1979; Wald 2004; Gold and Shadlen 2007; Roger Ratcliff and McKoon 2008; Urai et al. 2019; Gupta et al. 2023). Crucially, the decision-making module includes an urgency process that can initiate the movement, independently of the decision variable, thus accounting for the prevalence of express responses (Hernández-Navarro et al. 2021; Hawkins and Heathcote 2021) (Figure 1c). When the urgency process triggers the response, the targeted port of the initial trajectory is set by the sign of the decision variable at this moment *x_1_*. We refer to *x_1_* as the first read-out (Figure 5a, empty circle). Importantly, the vigor of the initial trajectory depends on the value of *x_1_*: the more evidence towards the selected port, the faster is the trajectory (Figure 5b). Following the same principle, when the decision variable hits one of the two decision bounds (*x_1_ =* ±**θ**), the initial response has always the maximum vigor. Regardless of which process triggers the initial response, due to sensorimotor delays, the latest sensory information is still in the processing pipeline at the moment of movement onset and thus does not affect the first read-out *x_1_*. The model makes a second read-out *x_2_* (Figure 5a, filled circles) once the whole stimulus evidence is fully integrated, allowing for the possibility to speed up or slow down the trajectory or even to reverse the decision (Figure 5c). Specifically, the trajectory is sped up if the evidence towards the initial choice increases with respect to the first read-out (i.e. |*x_2_|* > |*x_1_|*). When the evidence decreases (|*x_2_|* < |*x_1_|*), the initial trajectory is slowed down. When the second read-out is strongly at odds with the initial targeted port, a new ballistic trajectory is drawn towards the opposite port, implementing a change of mind (Figure 5a, bottom, yellow trace). Note that, while in the model we can identify all CoMs, only a subset of those will be classified as trajectory reversals using the detection method based on trajectory deflections as in rats. Trajectories were generated using the principle of minimum jerk which ensures smoothness, making the targeted port and the MT their only free parameters (Methods). Therefore, the updating of a trajectory, came down to drawing a new trajectory, that smoothly continued the initial one, but in which either the remaining MT was shortened or extended (vigor update), or the final target was switched (CoM).

**Figure 5.**
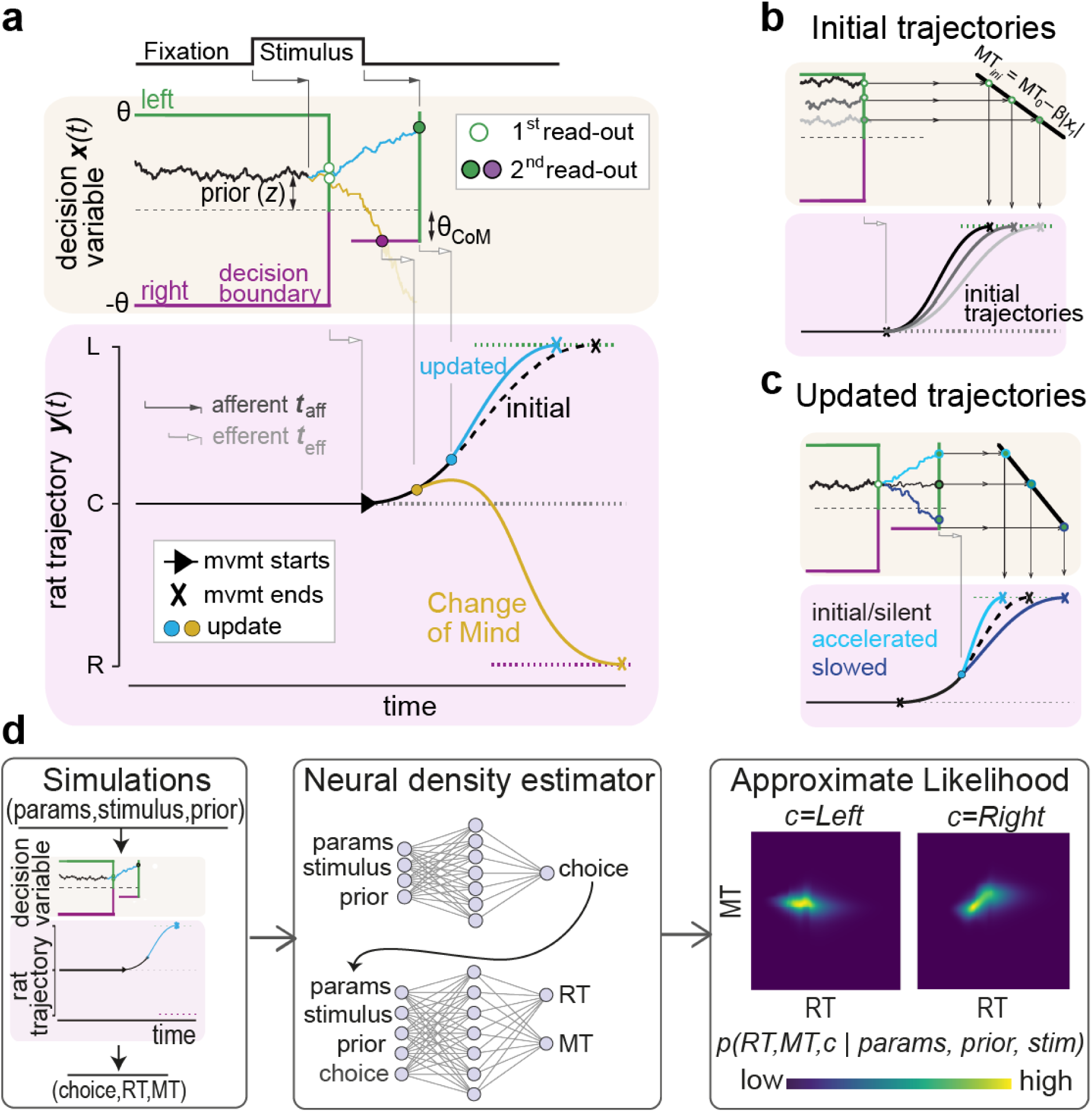
A joint statistical model for decision-making processes and response trajectories. **a)** The decision of the model is based on the decision variable *x*(t), which accumulates the sensory information towards the left (green upper bound) or right (purple lower bound) port following a drift-diffusion process. The value of *x*(t) at the trial onset is proportional to the prior evidence *z*. The stimulus is integrated after a delay with respect to the stimulus onset (afferent delay, t_aff_, gray solid arrows). Two example trials are shown in which the movement onset is triggered by a stimulus-independent process (Hernández-Navarro et al. 2021) that sets the time of the first read-out (vertical collapse of the decision boundaries). The first read-out, *x_1_*, triggers a leftward initial trajectory (bottom panel, black trace) with an efferent delay (*t*_eff_). In the standard trial (blue trace), sensory integration continues until the stimulus is fully processed (second vertical boundary): the second read-out *x_2_* (solid circles) updates the vigor of the trajectory by accelerating the movement (blue trajectory). Alternatively, the stimulus may contradict the evidence set by the prior (yellow trace), driving the decision variable into the change-of-mind bound (**θ**_CoM_, filled purple circle) and triggering a reversal of the trajectory (yellow trajectory). **b)** The movement time of the initial trajectory *MT_ini_* depends linearly on the absolute value of *x_1_*, where larger evidence converts into shorter MTs. **c)** The movement time is updated by a linear transformation of the second read-out: the trajectory is sped up if |*x*_2_| > |*x*_1_|, and slowed down otherwise. **d)** Model parameter estimation using Mixed Neural Likelihood Estimate. The model is simulated on 10 million trials, each with a different set of parameters and experimental conditions (stimulus and prior). Model simulations are then used to train an artificial neural network to approximate the conditional density of the data (i.e. choice, reaction time and movement time) given the model parameters and experimental conditions. Specifically, we train two different networks, that respectively learn *p*(*choice*|*parameters*, *prior*, *stimulus*) and *p*(*RT*, *MT*|*choice*, *parameters*, *prior*, *stimulus*). Finally, we estimate model parameters from experimental data using classical maximum likelihood estimation, but where the true (intractable) likelihood is replaced by the approximate likelihood provided by the trained network.

Fitting such a complex dynamical model is challenging because its likelihood cannot be expressed analytically. To solve this problem, we used Mixed Neural Likelihood Estimate (MNLE), a recently developed method to approximate the likelihood function of a statistical model using an artificial network (Boelts et al. 2022). In short, an artificial neural network was trained on 10 million simulations to approximate the joint likelihood of choice, reaction time and movement time, given the model parameters and the value of the prior evidence and stimulus strength.

The model captured very well the impact of the prior and stimulus evidence on the animals’ response trajectories (Figure 6; Supplementary Fig. 6). We fitted the 16 parameters of the model to match the choices, RTs and MTs of each rat individually. We then applied the same analyses used for the experimental data to the data generated from the fitted model. The model reproduces the experimental dependence on choice of prior, stimulus and RTs (Figure 6a-b) as well as the distributions of RT and MT (Supplementary Figs. 7 and 8). As the response vigor is determined by the value of the decision variable at both read-outs, the MT decreases as the prior and sensory evidence towards the response increases (Figure 6c-f). As in rats, model trajectories are faster compared to silent trials if the sensory evidence is congruent with the choice and slower if it is incongruent (Figure 6d). This follows directly from how the model updates the trajectories at the second read-out: stimuli tend to increase the initial evidence (with respect to silent trials) when they are congruent or decrease it when they are incongruent, causing acceleration or slowing, respectively. The model also reproduces the V-shaped dependence of the stimulus splitting time on RT (Figure 6g). The dependence is caused by a transition from fast responses (RT < *t_aff_+ t_eff_)* where the stimulus affects the second but not the first read-out, to longer responses (RT > *t_aff_+ t_eff_)* where it affects both read-outs (Supplementary Fig. 11a-c). Thus, the minimum splitting time provides an easily accessible upper bound for the sum of afferent and efferent times *t_aff_+ t_eff_* (Figures 2i and 6g and Supplementary Fig. 11d-f).

**Figure 6.**
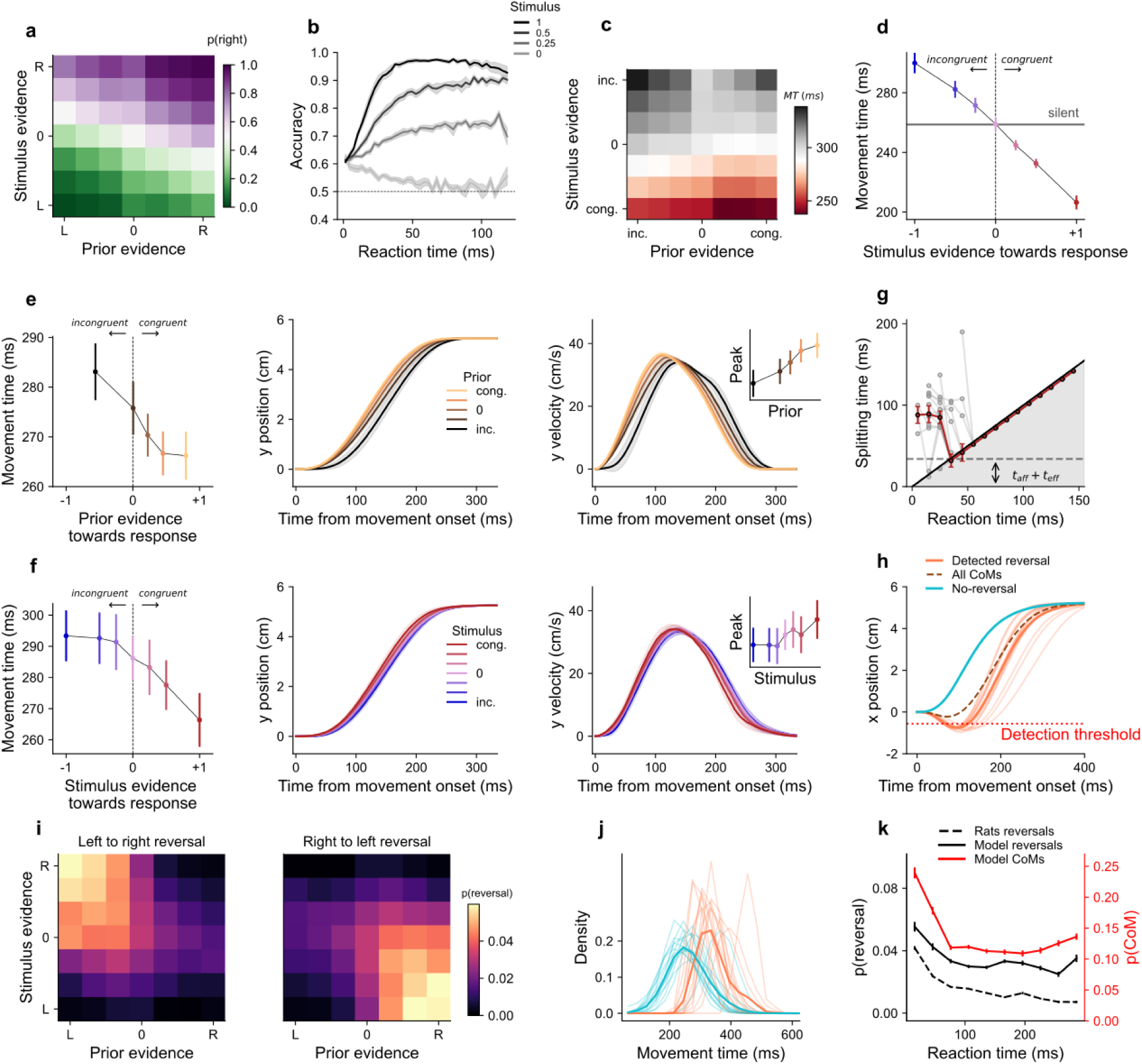
Simulations from the model fitted to individual rats replicate the key features of rat behavior. **a)** Proportion of right responses as a function of the stimulus and prior evidence (as in Figure 1b). **b)** Accuracy of the model as a function of RT, for different stimulus strengths (as in Figure 1d). **c)** Average movement time as a function of prior and stimulus evidence towards the selected port, averaged across animals (as in Figure 1h). **d)** Average movement time as a function of stimulus evidence supporting the decision, in express response trials (RT<50 ms) with prior evidence supporting the decision (prior evidence towards response z > median). Congruent stimuli accelerate responses w.r.t. silent trials (dashed line), while incongruent stimuli slow down responses, as in Figure 1i. **e)** Left: Median movement time in silent trials as a function of prior evidence towards the selected port (binned in 5 quantiles), averaged across animals. Error bars: s.e.m. Model trajectories (center) and velocities (right) in silent trials depending on prior congruence (as in Figure 2d, f). **f)** Median movement time (left), model trajectories (center) and velocities (right) in express response trials (RT< 50 ms) with small prior (|z| < 10% percentile), as a function of stimulus evidence towards the selected port. **g)** Splitting time for stimulus evidence as a function of RT, as in Figure 2i. **h)** Average simulated trajectories for CoM trajectories (dashed brown; including detected and non-detected trajectory reversals), detected reversals only (orange line), standard trajectories (blue line). **i)** Probability of reversals as a function of prior and stimulus evidence (see Figure 3i). **j)** Distribution of movement times for trajectories with changes of minds and standard trajectories. Legend as in Figure 3g. **k)** Proportion of trajectory reversals (black traces) as a function of reaction time for rats (dashed) and simulations (solid). Red line shows the proportion of changes-of-mind in the model.

Importantly, although the model was not fit to any property of the trajectories beyond their overall duration, it produces trajectory reversals in the same conditions as rats (Figure 6h). Specifically, these reversals occur mostly when the stimulus strongly contradicts the prior (Figure 6i, Supplementary Fig. 9). While changes of mind actually occur frequently, even when stimulus and prior weakly disagree, detection of trajectory reversals based on motion trajectories is more prominent for strongly disagreeing evidence (as strong prior leads to higher initial vigor, see details in Supplementary Fig. 12a-b). The majority of CoMs produced by the model occur early after movement onset, yielding kinks in the trajectories that are too small to be detected. Our model predicts that only around a quarter of the CoMs lead to detectable trajectory reversals (CoMs represented 15.67% of the total, vs 4.44% for detected reversals). Moreover, reversing trajectories incur a certain time cost, which explains why trajectories with reversals last on average longer than standard trajectories (Figure 6j). The model also explains why reversals are more frequent at short RTs (Figure 6k; see Figure 3j): while the first read-out reflects only the prior information, the second read-out also incorporates the sensory information, which can contradict and overcome the prior. By contrast, at longer RTs, the stimulus contributes to both read-outs, making changes of mind less frequent (Supplementary Fig. 12c). Changes of mind in the model are intimately related to the existence of proactive responses: as we gradually change the model parameters to extinguish the fraction of proactive responses, the proportion of CoM also vanishes (Supplementary Fig. 11g). Overall, there is a compelling fit between our model and the experimental data, in terms of movement time, the timing of the impact of prior and sensory evidence onto trajectories, and the trajectory reversals.

The alignment between the data and our model supports the hypothesis that the evidence-accumulation process determines motor response kinematics at two distinct time points. What is the distinctive impact of these two time points? To answer this question, we evaluated three alternative models where we removed or altered either of the two read-outs: a model where the first read-out was replaced by a random initial choice (*random initial choice model*, Figure 7b); a model where the second read-out was removed (*no trajectory update model*, Figure 7c); and a model where the second read-out only occurs when reaching the CoM boundary (i.e. trajectories were not updated if the response was confirmed, *no vigor update model,* Figure 7d). Importantly, each of these alternative models failed to capture important characteristics of movement time and/or trajectory reversals found in animal behavior. The *random initial choice model* produces by construction 50% of CoMs, disrupting the dependence of CoMs on the decision variables (prior and stimulus evidence; Figure 7b, iii). In this model, unlike in experimental data, reversals are more frequent when both the prior and the stimulus evidence are weak (Figure 7b, iv), so that the weak vigor leads to a change in direction that is slow enough to cross the reversal detection threshold. By construction also, the *no trajectory update model* never produces trajectory reversals (Figure 7c, iv). Moreover, movement time in express responses of this model are largely decoupled from the stimulus evidence (Figure 7c, ii), as the trajectory is triggered at the first read-out before the stimulus impacts the decision variable (Figure 7c, i). For the same reason, the decoupling between stimulus and MT in express responses is also observed in the *no vigor update model* (Figure 7d, ii), although this model produced CoMs and reversals exactly as the full model (Figure 7d, iii-iv). These results evidence the existence of at least two read-outs impacting the motor response kinematics that, based on the evidence accumulated, accelerate, slow down or reverse the trajectory of the rat.

**Figure 7.**
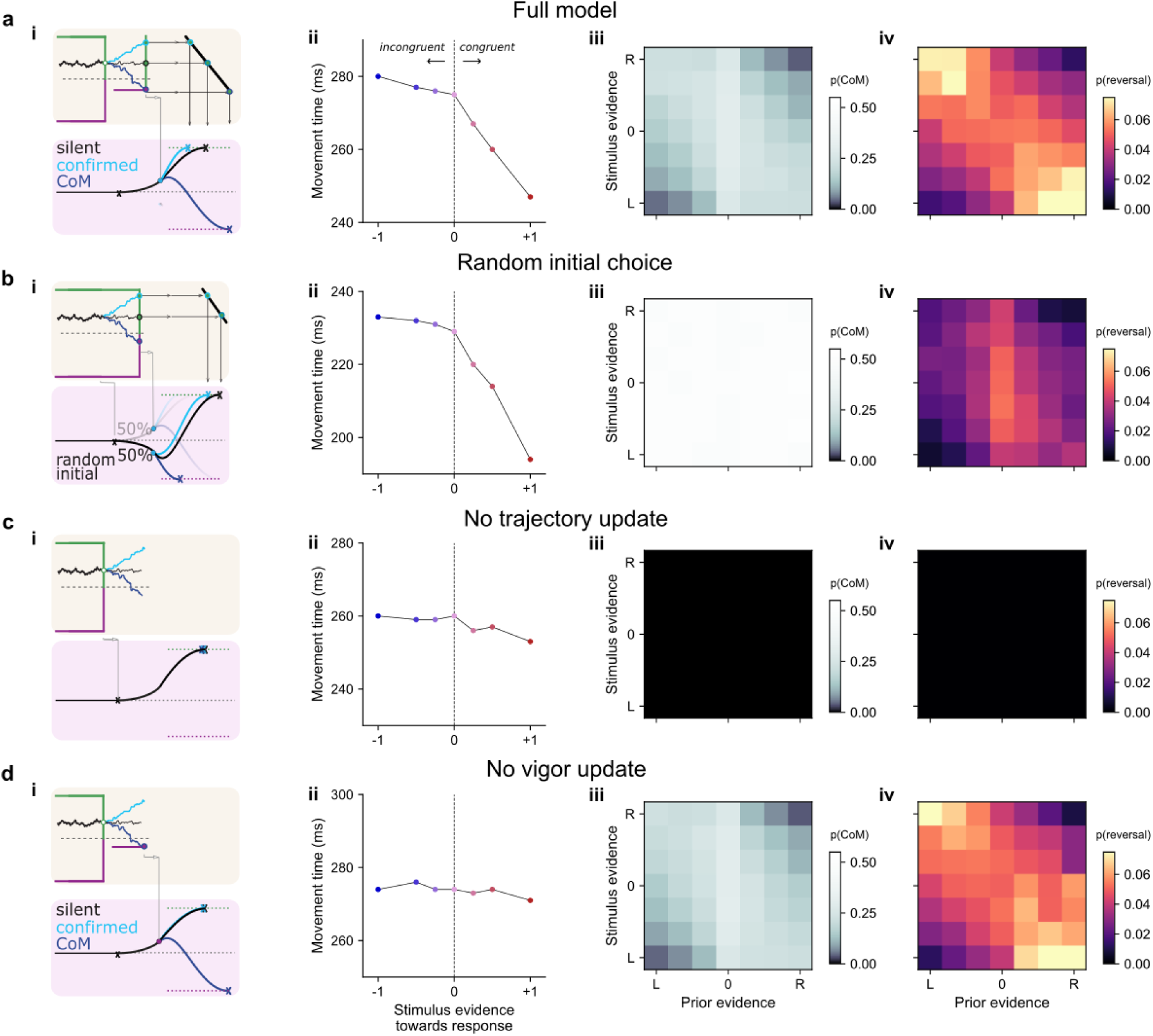
Comparison between alternative models: full model (**a**), model with random initial choice (**b**), model with no trajectory update (**c**) and model with no vigor update (**d**). **i)** Cartoon illustrating each model. **ii)** Median movement time in express response trials (RT< 50 ms) with small prior (|z| < 10% percentile), as a function of stimulus evidence towards the selected port. **iii-iv)** Probability of CoM (iii) and reversals (iv) as a function of prior and stimulus evidence.

## Discussion

The advent of automatic techniques to extract the position of body parts from video footage has opened the path to rich quantitative analyses of motor behavior in freely moving animals. However these techniques had never been deployed in the context of decision-making tasks in mammals, (except for (Kane, Senne, and Scott 2023)). Through thorough analysis, we demonstrate that the response trajectories observed in rats performing a two-alternative forced choice task can be encapsulated by a dynamical model in which response movement is tightly steered by the evidence-accumulation process. Our model captures the dependence of trajectory reversals on stimulus, prior and reaction time, even though it was not fitted to any characteristics of the trajectory other than its overall duration. The same model successfully replicated the response trajectories described by human subjects performing an analogous task on a tactile screen. While most previous theoretical work has focussed either on decision-making processes before a response is initiated or on the unfolding of reaching movements (Wispinski, Gallivan, and Chapman 2020), our joint modeling of decisions and response trajectories provides a key step to dig further into the interactions between decision-making and motor control (Friedman, Brown, and Finkbeiner 2013; Lepora and Pezzulo 2015). Since motor-related activity has a large and global impact on neural activity in awake mice (Stringer et al. 2019; Musall et al. 2019), providing a foundation for models of joint decision-making and decision-related movements can significantly help the analysis and understanding of neural data.

### Accumulated decision evidence impacts directly and gradually motor vigor

The modern view of decision-motor interaction encompasses parallel processing circuits that can decide while moving and move while deciding (Gordon et al. 2021). In line with this perspective, our work shows a direct and graded relationship between the accumulated decision evidence in a perceptual task and the vigor with which rats execute their response movements. To the best of our knowledge this is the first time that the vigor of a complex body movement has been tightly linked to the decision making process. Previous studies have either not found any relationship of MT with e.g. stimulus evidence (Zariwala et al. 2013; Lak et al. 2014) or have used paradigms that conflated both RT and MT (by reporting the time from stimulus onset to movement offset), which hindered the possibility of isolating the impact of accumulated evidence on movement vigor (Kane, Senne, and Scott 2023). Saccadic eye movements (either response related or not) are faster when perceptual evidence supporting the choice is larger (Seideman, Stanford, and Salinas 2018; Korbisch et al. 2022). Here we show that the link also exists for complex movements involving the full body, and describe in detail the relationship between the dynamics of the decision variable and the kinematics of the movement. More broadly, because response vigor is also modulated by the intrinsic action value (Milstein and Dorris 2007; Xu-Wilson, Zee, and Shadmehr 2009; Reppert et al. 2015; Summerside, Shadmehr, and Ahmed 2018), we can construe that vigor is based on the expected reward from the corresponding action, which depends on the probability of obtaining the reward and which can ultimately be approximated by the decision variable.

Previous studies have identified trajectory updates in scenarios where the later part of the stimulus contradicts the earlier part (Resulaj et al. 2009; van den Berg et al. 2016) or when salient sensory evidence is contradicted by information that requires top-down processing (Nakahashi and Cisek 2023), making the subjects reverse their initial trajectory and reach the opposite option, i.e. perform a change of mind. We found that changes of mind can also occur when the stimulus violates a prior expectation, a situation that happens frequently both in naturalistic conditions (e.g. a goalkeeper during a penalty) and in the lab (e.g. the Posner task (Posner 1980)). The existence of such prior-based changes of mind is directly dependent on the prominence of proactive responses (Supplementary Fig. 11g). When proactive responses were removed from the model, changes of mind solely driven by the stimulus (Resulaj et al. 2009; van den Berg et al. 2016) were absent as fluctuations in sensory evidence were not large enough to drive the decision evidence from one boundary to the CoM boundary. More fundamentally, beyond the difference in the origin of CoMs, our results suggest that trajectory updates occur routinely: even when the final target is not changed, newly acquired sensory information always leads to a slowing or speeding of the response plan (Figures 2e,g and 4f). Importantly, this adjustment is graded as rats accelerate when the newly acquired sensory information confirms their initial choice and slow down when it contradicts it. The analysis of trajectories unveiled the latency of this update: the time elapsed between stimulus onset and the visible deflection of the trajectory, which encompasses a whole range of delays, from sensory processing to motor activations, can be as small as 60 ms. We verified in further model simulations that this minimal delay was composed by the sum of the afferent delay, the efferent delay, and a residual time required to detect a statistically significant effect (Supplementary Fig. 11 a-c). In humans, we found a minimal delay on the order of 200 ms (Smeets, Oostwoud Wijdenes, and Brenner 2016; Nakahashi and Cisek 2023), which matches the delay necessary for interrupting a response movement (Schultze-Kraft et al. 2016); and is much larger than the 30 ms period required for minimal sensory integration (Stanford et al. 2010) which makes up for only one link of the sensorimotor chain.

### Statistical modeling of response trajectories

Accumulation-to-bound models such as the Drift-Diffusion Model (DDM) (Roger Ratcliff and McKoon 2008) describe the pattern of choices and reaction times (R. Ratcliff 1985; Palmer, Huk, and Shadlen 2005; Bogacz et al. 2010; Pardo-Vazquez et al. 2019), but they do not capture the impact of the decision-making process on the motor trajectories (although see (Kiani, Corthell, and Shadlen 2014)). Our study showcases how extending the quantitative toolkit of decision processes to model response trajectories can illuminate the link between decision-making and motor processes. Although the actual response involves the coordination of many effectors to reach the selected port, we found that the trajectories of the rat snout were usually smooth and stereotypical, making them amenable to standard statistical modeling.

The model makes several predictions for further experiments. First, it predicts that our subjects change their mind much more often than what is apparent from the identified trajectory reversals. These latent changes of mind are akin to the “neural reversals”, i.e. changes in the sign of the decision variable driven by changes in the stimulus evidence, as observed in rats (Boyd-Meredith et al. 2022) and monkeys (Kiani et al. 2014; Kaufman et al. 2015; Peixoto et al. 2021). Second, consistent with previous work in humans (Friedman, Brown, and Finkbeiner 2013), our model includes a second read-out of the decision variable after the movement is initiated to correct or adjust the initial response. The actions occurring between read-outs are thus ballistic *submovements* conveniently concatenated to minimize sharp changes in velocity. One open question is whether more updates could occur along the trajectory (Friedman, Brown, and Finkbeiner 2013; Lepora and Pezzulo 2015) the brain could continuously transform decision variables into changes in the response trajectory (Friedman, Brown, and Finkbeiner 2013; Lepora and Pezzulo 2015). The implementation of such a continuous model would require accommodating the dynamics of evidence accumulation, which typically exhibit fast sudden changes in the decision variable, to the constraints of optimal motor control which imposes smooth, gradual changes in velocities and acceleration. Our discrete-updating model may be an intermediate solution by which trajectories are constrained by the movement kinematics but are tightly influenced by the evolution of decision variables across time.

Previous work in humans has found that trial history effects in perceptual decisions can be mediated by a bias in the drift of the accumulated evidence (Urai et al. 2019). We however modeled the influence of prior evidence purely as an offset in the initial value of the decision variable (Figure 5a), since the impact of prior on rat choices and vigor was strong for express responses (Figures 1d-e and 2d,f) and decreased with RT (Supplementary Fig. 1d). Moreover, the position of the rats’ snout depended on the prior evidence already during fixation, before sensory integration started. The difference between our data and the mentioned study may be due to a different origin of the trial history effects. Moreover, the model did not incorporate idiosyncratic asymmetries in the integration of prior and stimulus, observed in some rats and humans (see Supplementary Fig. 2 and Figure 4l, respectively). Introducing fixed biases within the prior evidence as well as in the evidence accumulation drift could possibly account for these individual preferences. Another aspect that our model ignores is the reciprocal effect that motor kinematics can have on the decision making process. In other words, on top of the impact of the accumulated evidence on the motor trajectories, the motor components may very well impact the decision (Cos, Bélanger, and Cisek 2011; Lepora and Pezzulo 2015; Shadmehr, Huang, and Ahmed 2016; Kane, Senne, and Scott 2023), a phenomenon known as embodied decision-making. For example, approaching one port may lead to a downweighting of contradictory sensory evidence (or an increase in the change-of-mind boundary), in order to avoid a late and costly reversal. Such a phenomenon is closely related to the confirmation bias by which subjects discard evidence contradicting an initial choice (Nickerson 1998). Note however that the boundary for changes of mind was close to null in most animals (Supplementary Fig. 10), unlike in humans (Resulaj et al. 2009), indicating a lack of bias to confirm their response even as they already departed towards the associated port.

### Making decisions in a continuous world

The similarity between the behavior of rats and humans in our task suggests that the strategy used by both species to adjust movements endpoints and vigor while processing asynchronous information evolved before the two species diverged around 75 million years ago (Striedter and Glenn Northcutt 2019). Our results also suggest that these same mechanisms control very different types of motor response, such as the full-body orientation or the movement of a finger. Consistent with this hypothesis, the basal ganglia, which regulates motor vigor (Turner and Desmurget 2010), is a highly preserved area since its emergence in the first vertebrates 500 million years ago (Grillner and Robertson 2016). This conservation may reflect the fact that embodied decisions, in which the subject acts while deciding and decides while acting (Gordon et al. 2021), have been crucial for survival throughout the evolutionary history of vertebrates. Beyond their similarities, humans differed from rats as they performed many more reversals. Part of this difference may be due to the lower energetic cost of a change of trajectory in the finger compared to full-body responses. Indeed, many human reversals occurred when the finger was already close to the target port initially selected. Future research should explore the conditions that lead to reversals when the time-varying stream of sensory information continues throughout movement execution (Kane, Senne, and Scott 2023).

## Methods

### Rat behavioral protocol

15 male Long-Evans rats performed a free reaction time (RT) two-alternative forced choice (2AFC) intensity level discrimination (ILD) task (Pardo-Vazquez et al. 2019). We summarize the structure of the task below, for further details see (Hermoso-Mendizabal et al. 2020; Hernández-Navarro et al. 2021). Rats initiated a trial by maintaining fixation at the center port during 300 ms. A broadband noise stimulus was then presented at two speakers, and lasted until the rat retracted from the port and initiated a response. The two lateral ports were only about 5 cm from the central port, which allowed rats to rapidly execute their response trajectories by only moving their upper body (see Supplementary Video 1). The relative stimulus strength (i.e. the average difference in sound intensity between the two speakers) changed every trial between different values: 0 (mean equal intensity), 0.25, 0.5 and 1 (stimulus played on the correct side only). This resulted in 7 values of stimulus evidence (i.e. stimulus strength signed by the stimulus category), ranging from −1 (clear evidence to the left) to +1 (clear evidence to the right). The interaural level difference fluctuated dynamically within each stimulus along the predetermined mean. Correct responses were rewarded with 25 μl of water and incorrect ones punished with a 2 s timeout. The stimulus category (left vs right dominant) was drawn based on the current block: in repeating blocks, the stimulus category was repeated with probability P_rep_=0.8; in alternating blocks, the stimulus category was repeated with probability P_rep_= 0.2. In 6 out of 15 rats we also introduced a small fraction of silent trials in which, despite not presenting any stimulus, rats elicited a valid response (mean±SD 6.7 ± 0.6% of all trials; Group 2 in Table 1). Trials with reaction times above 400 ms or motor time above 1000 ms were excluded. The behavioral setup was controlled by BPod, an open control system for precision animal behavior measurement (by Sanworks) and the task was run using the Python-based open software package PyBPod (http://pybpod.com/).

**Table 1.** Rat groups.

| Animal groups | # rats | # sessions<br>(mean±std) | # trials per rat<br>(mean ± std) | # silent trials<br>per rat<br>(mean ± std) |
| --- | --- | --- | --- | --- |
| Group #1 (ILD task):<br>LE36, LE37, LE38, LE39, LE40,<br>LE41, LE84, LE85, LE86 | 9 | 131 ± 45 | 59,870<br>± 25,933 | 0 |
| Group #2 (ILD task + silent<br>trials): LE42, LE43, LE44, LE45,<br>LE46, LE47 | 6 | 283 ± 45 | 123,983<br>± 26581 | 8,411<br>± 2,352 |
| TOTAL | 15 | 2,889 | 1,282,733 | 50,468 |

### Model of prior evidence

We defined the prior evidence *z* as the magnitude in each trial of the choice bias caused by the recent history of trials. As we described in detail in a previous work (Hermoso-Mendizabal et al. 2020), rats leverage the trial sequence statistics to develop a prior expectation on the next rewarded side. Because history-dependent biases disappeared following an error (i.e. the prior did not guide the decision) (Hermoso-Mendizabal et al. 2020), in the remainder of the analyses we only analyzed trials following correct responses (77.13 ± 2.65 %). This prior expectation can be recapitulated by using a Generalized Linear Model of choices which quantifies the weight of different elements of recent history on choices (Busse et al. 2011; Frund, Wichmann, and Macke 2014; Abrahamyan et al. 2016; Braun, Urai, and Donner 2018; Hermoso-Mendizabal et al. 2020). In rats, we used the same logistic regression model as in (Molano-Mazón et al. 2023) - see the reference for full model description. The model regressors include the previous repetitions/alternations (with different weights depending on whether each of the responses in the transition were rewarded) and previous rewarded and non-rewarded responses occurring in the last 10 trials. The model also includes a regressor for current stimulus *S_t_* (defined as the average intensity difference between the two tone sounds) and a fixed side bias. After fitting the GLM model to the choices of each individual animal, the prior evidence *z* was defined in each trial as the sum of the trial-history regressors weighted by their corresponding regression weights. For humans, due to the lower number of trials, we used an exponential kernel as a basis function for the impact of the different history regressors (Supplementary Fig. 3), as (Frund, Wichmann, and Macke 2014).

### Automatic tracking of rat position

A USB camera OV2710 CMOS (3.6 mm lens, 30 to 120 frames per second, 640 x 480 pixels per frame) was placed at approximately 30 cm from the floor level, recording the animal movements from above. The camera had no infra-red filter, so the rats could be recorded without being disturbed by visible light. Snout coordinates in the horizontal plane were extracted from videos using DeepLabCut (v1.11 using a resnet50 architecture) (Mathis et al. 2018). To train the network to extract the rats pose, a total of 480 images from several subjects were manually labeled. We used nine points in the rats body: the base of tail, a point on the bottom of the back, the midpoint between the two omoplates, the tip of the two ears, the two eyes, a point at the snout base and a point at the tip of the snout (Supplementary Video 1). We then used the point at the snout base to extract response trajectories as the tip of the snout was often occluded when rates poked into the ports. 95% of the images were used to train the model, while the remaining 5% was used to evaluate its performance. The deviation (RMSE) of the snout tracking done by the model from the ground truth (human-labeled images) was 3.71 pixels for the test set (1.04 pixels for the training set). Each pixel corresponded to 0.07 centimeters.

### Preprocessing of the trajectories

The output of this processing pipeline was a time-series for the coordinates of the animal snout in two dimensions *(x,y).* To compensate for small camera movements across sessions, pixel space was re-referenced on every session such that the central port was located at *y=0* and lateral ports at coordinates *y=-±75* pixels (*y=±5.25 cm)*. The time-series were interpolated linearly to infer trajectories at a time resolution of 1 ms. We excluded all trajectories for which the detected *y* position was above +100 pixels or below −100 pixels (+7 cm or −7 cm, respectively) at any time during the orienting trajectory (1.12 % of trajectories), or when the end point of the trajectory was inconsistent with the registered response port (0.41%). All trajectories were re-referenced temporally to motion onset (*t=0*), i.e. the time at which the animal left the central port.

### Average trajectories

We obtained trial-averaged trajectory traces and velocity traces for different values of the stimulus and prior evidence. We first flipped the sign of trajectories ending at the right port so that all trajectories end at positive values. We padded the coordinates of the snout at constant value after reaching the response port. Further, on each trial, we shifted all coordinates along the y-coordinate so that *y=*0 corresponds to the average position of the animal in the last 100 ms before motor onset. The stimulus evidence towards the response corresponds to the stimulus strength (between 0 and 1) signed by the congruence with the final response. For example, if the stimulus category is rightward but the selected port is leftward, the stimulus is incongruent with the response, and so the stimulus evidence *towards the response* is negative. The prior evidence towards the response is defined similarly. The signed prior evidence was binned into 5 quantiles. Instantaneous *y* velocities were computed by finite differences, i.e. they corresponded to the difference of *y* coordinates in successive frames divided by the time resolution.

### Movement time analysis

Movement time was defined as the duration between central port withdrawal to lateral port entrance, both registered by photogates at the corresponding ports. We measured the raw impact of prior evidence on MT by focussing on silent trials, to avoid the confounding effect of sensory evidence. We measured the raw impact of stimulus evidence on MT by focussing on trials with small prior (|z|<10% percentile), to reduce the confounding effect of prior evidence and on express responses (RT<50 ms) in which RTs are independent of stimulus evidence, in order to reduce a possible confounding indirect impact of stimulus on MT reaction time and then on MT). Finally, to test whether sensory evidence can speed up and/or slow down trajectories, we focussed our analysis on trials whereby the prior strongly supported the final decision (prior evidence towards response z > median) in order to minimize the proportion of reversals; and in express response trials, to reduce the impact of stimulus evidence on RT. We also performed a linear regression analysis for MT in each animal separately, using three different regressors: the stimulus strength and the prior evidence towards the responses; and the index of the trial within the session, an indicator of the animal satiety and fatigue (Hernández-Navarro et al. 2021). All regressors were z-scored.

### Splitting time

To estimate the earliest time at which stimulus information impacted the trajectories, we computed the splitting time of trajectories associated with different stimulus strengths. Here, we first flipped the sign of the *y* coordinates when the stimulus category was rightwards (independently from the actual response), so that positive *y* values correspond to trajectories towards the correct port. For each reaction time bin (bin size=10 ms), we computed the Pearson correlation across trials between the position of the animal along the *y* axis and the stimulus strength. We extracted the time-series of p-value for these correlations and identified the splitting time as the last non-significant time point using a criterion of p=0.05. This analysis was performed separately for each animal.

### Trajectory reversals

For each trial, we measured the maximal position of the animal along the mean-shifted *y* axis (see Average Trajectories) in the direction contrary to the response port and labeled this as the reversal point. Trials were identified as reversal trials if this value exceeded a certain detection threshold (8 pixels, i.e. 0.56 cm). We applied more stringent criteria (i.e. higher detection threshold) which yielded a smaller fraction of CoMs. However, qualitatively, results were unchanged when using a more stringent detection threshold.

### Statistical model of decision-making and response trajectories

The generative model is composed of two modules: a decision-making module, which integrates both the prior based on the history of previous trials and the stimulus evidence from the current trial, and a motor module that translates the accumulated evidence into a smooth orienting trajectory towards the corresponding port. The overall principle of the model is to jointly maximize the probability for correct choices and minimize movement costs (Rigoux and Guigon 2012; Lepora and Pezzulo 2015). The decision-making module is based on the architecture of the Parallel Sensory Integration and Action Model (PSIAM) (Hernández-Navarro et al. 2021). In the PSIAM, a response is initiated wherever either of two parallel processes first reaches a boundary: an Evidence Accumulation process that integrates decision evidence over time and initiates the response when it reaches a bound, akin to the Drift Diffusion Model (Roger Ratcliff and McKoon 2008); and an Action Initiation process that can trigger the initiation of the response proactively, i.e. independently of the decision evidence integration, at a timing determined by a stochastic process. The Evidence Accumulation process is formalized as a decision variable x(t) that represents the relative evidence in favor of each choice, with positive and negative values supporting the leftward and rightward choices, respectively. Its initial value (at fixation onset) is equal to the prior evidence *z* multiplied by the model parameter *z_p_*. The dynamics of the decision variable x(t) are subject to leak λ, and integrate the stimulus evidence *s(t)* with drift *a_P_* and an afferent delay *t_aff_*, following:

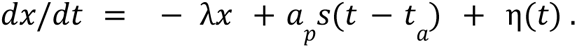

where η(*t*) is a white noise process of variance 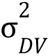. If the decision variable reaches one of two symmetric decision bounds ± **θ**_DV_, the model selects the corresponding response (first read-out) and the trajectory towards the selected port is initiated. A response can be also initiated if the Action Initiation process reaches its associated boundary first. This process is an independent drift-diffusion process with diffusion coefficient 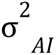, a single boundary (θ*_AI_*) and drift that depends linearly on trial index as *a_AI_* = *v_AI_* + *w_AI_ TrialIndex*, where *v_AI_* and *w_AI_* are the intercept and slope, respectively. The Action Initiation variable starts integrating after some delay (*t_AI_*) from the fixation onset. In the case that the response is triggered by the Action Initiation, the side of the response is determined by the sign of the decision variable at that time *x*_1_ (1st read-out). In either case, the initial response is started after the first read-out with an efferent delay *t_eff_*, which triggers the offset of the stimulus. Evidence integration continues after this first read-out and until the stimulus is fully integrated (i.e. for a further delay equal to the sum of afferent and efferent delays *t_eff_* + *t_aff_*). If during this interval the decision variable hits the boundary for changes of mind (set at a value *θ_COM_* with the sign opposite to the initial response), the initial response is reversed (Figure 5a, yellow traces). Otherwise, the trajectory is maintained, and the second read-out (i.e. value *x*_2_ of the decision variable after a delay of *t_eff_* + *t_aff_* with respect to the first read-out) only modulates the vigor of the rest of the trajectory towards the response port (see below).

The motor module translates the decisions onto motor trajectories. The initial trajectory is computed by first selecting the initial Movement Time, which depends linearly on the absolute value of the evidence accumulated at the first read-out |*x*_1_| and on the trial index:

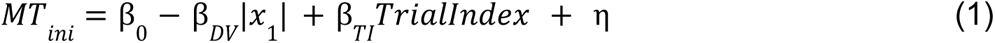

where η is drawn from a Gumbel distribution of variance 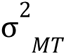 and mode at zero (this distribution was used to produce right-skewed MT distribution as observed in animals, see Figure 3h). This initial MT is then converted into a full trajectory by applying the principle of minimal jerk (Flash and Hogan 1985), which is known to produce good approximations to ballistic movements. The principle specifies the full trajectory *y(t)* between a starting point *y_0_* and an endpoint *y_E_* given the movement duration (or MT), and the velocity and acceleration at the start point and endpoint. This trajectory minimizes the overall jerk, that is the integral of the square of the third-order derivative of position over the entire interval. The solution is given by a 5th order polynomial: 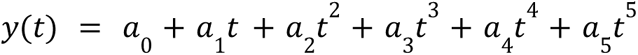 (Shadmehr and Wise 2004). The vector of coefficients ***a*** is found by solving a linear system specifying the boundary conditions on *y*(*t*), *y*’(*t*) and *y*’’(*t*) at the initial and final points *t*_0_ and *t_E_*:

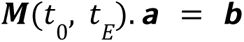

where

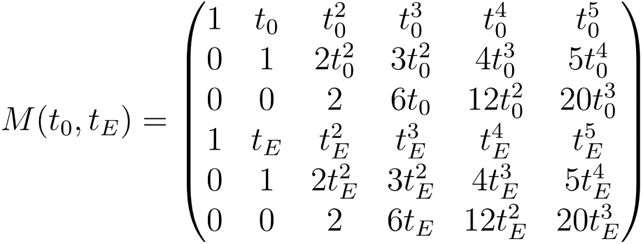

and the vector *b* is determined by the boundary conditions 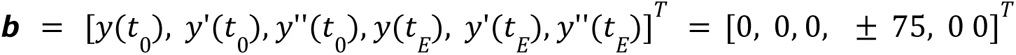. Note that we assume null velocity and acceleration along the y-axis at the initial and ending point (the sign of the position at *t_E_* depends on the selected response port).

Inverting this system gives ***a*** = ***M***(*t*_0_, *t_E_*)^−1^***b***. Imposing *t*_*o*_ = 0 for the initial trajectory,

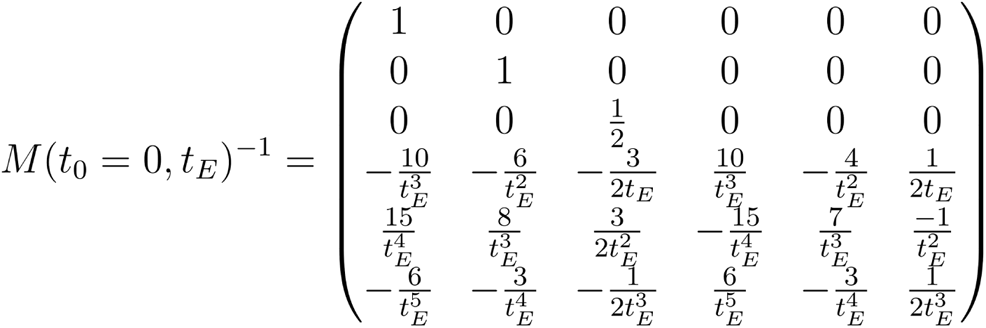

with *t_E_* = *MT_ini_*. The initial trajectory is later updated following the second read-out at time *t_U_* = *t*_2_ + *t_eff_*, with *t*_2_ being the time of the second read-out. If that second read-out leads to a change of mind, the endpoint is changed to the alternative response port (Friedman, Brown, and Finkbeiner 2013). If no change of mind is produced, the response port stays unchanged but the end time is updated. In all cases, the endtime changes according to

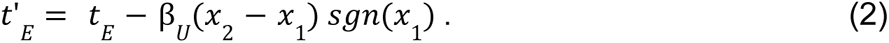

In other words, the trajectory is sped up w.r.t the original trajectory if the decision variable at the second read-out is larger in amplitude than at the first read-out, and is slowed down otherwise. The updated trajectory between *t_U_* and *t*’*_E_* is drawn again using the minimum-jerk principle, solving a new linear equation as seen above but where the initial conditions correspond to the position, velocity and acceleration of the initial trajectory at time *t_U_* (due to the continuity of these variables). Model simulations produced trajectories that we sampled at a 1000-Hz rate.

The model contains 18 parameters overall: 5 parameters related for the Action Initiation process (*t_AI_*, *v_AI_*, *w_AI_*, σ*_AI_*, θ*_AI_*), 8 parameters for the Evidence Accumulation process (*t_aff_*, *t_eff_*, *Z_P_*, *a_p_*, λ, σ*_DV_*, θ*_DV_*, θ*_COM_*) and 5 parameters for the response trajectories (β*_o_*, σ*_MT_*, β*_DV_*, β*_TI_*, β*_U_*). Parameters σ and σ were set to one to avoid identifiability issues for the AI and DV process.

### Alternative models

The following alternative models were simulated, with parameters fitted from the full model. In the *random initial choice model,* there is no reading of the decision variable in the first read-out. Therefore, the initial choice is randomly selected between left and right with 50% probability, and the initial trajectory is independent of the accumulated evidence (i.e. β*_DV_* = 0, so *MT_ini_* = β_0_ + β*_TI_TrialIndex* + η from Equation 1). A single read-out is performed after all evidence has been integrated, based on vertical boundaries. The updated movement time is computed using the absolute value of the decision variable at that instant. CoMs emerge when the first (random) and final choice are at odds, which by construction occurs in 50% of trials. In the *no trajectory update model,* the first read-out occurs as in the full model, but there is no update at all (i.e. β*_U_* = 0, so *t*’*_E_* =*t_E_* from Equation 2). This model does not produce CoMs, since there is no second reading of the decision variable. Finally, the *no vigor update model* is identical to the full model except that the trajectory is only updated in the case the DV hits the CoMs boundary (i.e. β*_U_* ≠ 0 when a CoM occurs, β*_U_* = 0 otherwise). This model is conceptually similar to the change-of-mind model in (Resulaj et al. 2009).

### Model fitting

The model parameters were fitted to the joint pattern of choices, reaction times and movement times, separately for each rat. Because the likelihood of such a model cannot be expressed analytically, we used Mixed Neural Likelihood Estimator (MNLE), a simulation-based method to approximate the likelihood of a model designed for mixed data, as typically found in decision-making tasks (Boelts et al. 2022). Here, the mixed data corresponds to one binary variable (choice) and two continuous variables (RT and MT). MNLE consists of training an artificial neural network that takes model parameters as input and outputs a joint probability distribution for the dependent variables. We extended the method to also include per-trial conditions (stimulus evidence *s_t_*, prior evidence *z_t_*, trial index *ti_t_*) as extra input to the network. Thus the method approximated *p*(*c*, *RT*, *MT*|θ, *s_t_*, *Z_t_*, *ti_t_*). More precisely, the method relies on training two separate networks (Figure 5d), one that learns the conditional probability *p*(*c*|θ, *s_t_*, *Z_t_*, *ti_t_*), while the other learns the conditional density *p*(*RT*, *MT*|*c*, θ, *s_t_*, *Z_t_*, *ti_t_*). The network was trained on 10 million simulations of the original model: in each simulation, the model parameters were sampled from uniform distributions over plausible values, while the per-trial conditions corresponded to one trial randomly selected from the rat experimental dataset.

We confirmed that the trained network provided a good approximation to the true likelihood in three different ways. First, we compared its output for a fixed parameter set to the density obtained from 500.000 model simulations at the same parameter values (Supplementary Fig. 4). This was repeated for a set of 6 trial examples covering all prototypic configurations of prior-stimulus association and different levels of trial index. Second, we checked the robustness of the method by comparing the approximated likelihood of two networks trained with two different sets of 10 million simulations (Supplementary Fig. 5a-f). Finally, we confirmed that the likelihood approximation improves as the size of the training set increases (Supplementary Fig. 5g). We then fitted the model for each individual rat through maximum likelihood estimation, using the approximate likelihood provided by the trained network. To limit the impact of possible outliers on parameter estimation, we used a mixture of two distributions, the approximated distribution and a contaminant distribution (Hernández-Navarro et al. 2021):

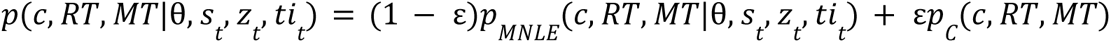

The contaminant was simply set as a uniform distribution (with RT bounded between 0 and *RT_max_*=1, and MT bounded between 0 and *MT_max_*=0.4), i.e. *p_C_*(*c*, *RT*, *MT*) = 1/(2*RT_max_MT_max_*). The mixture parameter *ε* was set to 10^-3^. Because fixation breaks (i.e RT<0) have no associated choice nor movement time (the trial was aborted), we replaced the joint likelihood term for these trials by the likelihood of the reaction time, i.e. we defined the overall likelihood as 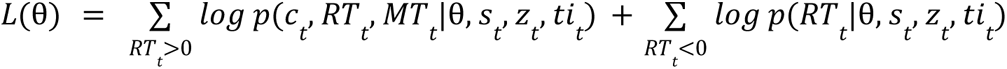. The likelihood of the reaction time for these fixation breaks can be expressed analytically as the RT distribution in negative RTs (which corresponds only to AI-triggered responses) follows an inverse Gaussian distribution (Hernández-Navarro et al. 2021). The optimization procedure relied on Bayesian adaptive direct search (BADS) (Acerbi and Ma, 2017), implemented in the PyBads package.

### Human behavioral paradigm

We tested *n*=20 volunteer participants (10 females and 10 males, 19 right-handed and 1 left-handed, age 19-26) performing a task that mimicked the rodent task. The task was performed on an iPad tablet with resolution 2048 x 1536 pixels (19.7 x 14.8 cm), with 60 Hz frame rate at an approximate viewing distance of 50 cm, using the StimuliApp software (Marin-Campos et al. 2021). In each trial, a black screen displayed three gray circles: the starting point at the bottom of the screen, and two target points positioned symmetrically. The target points were located at coordinates (−600, 600) and (600, 600) pixels, corresponding to (−5.76, 5.76) and (5.76, 5.76) cm, respectively, in relation to the starting point. Subjects initiated each trial by maintaining the index finger of their preferred hand pressed against the starting point for 500 ms (fixation period). The white-noise stimulus was then presented binaurally using Sennheiser headphones. The relative interaural level difference of the stimulus changed every trial between different values: 0 (mean equal intensity), 0.1, 0.2 and 0.4; the mean intensity was constant throughout the stimulus. Subjects had to slide the finger towards the target point at the side corresponding to the loudest stimulus. The stimulus was interrupted as soon as the finger slid away from the starting point. Feedback was provided after reaching the target point. Correct and incorrect choices were indicated by the whole screen turning green and red, respectively, for 300 ms. Importantly, subjects had to start moving the finger within 300 ms after stimulus onset, otherwise the trial was considered a missed trial and the screen turned yellow. This time constraint introduced an urgency component which nudged participants into relying more on prior expectations. The screen also turned to yellow if subjects initiated a response before the end of the fixation period. The trial sequence statistics replicated the task used for the rats. The probability of repeating the previous stimulus category (left/right), P_rep_, varied between Repeating blocks (P_rep_=0.8) and Alternating blocks (P_rep_=0.2). Blocks changed every 80 trials. Subjects were not informed about this correlation structure. They performed a training of 200 trials before completing the task. The session lasted on average 1 hour and 40 minutes (interleaved by pauses every 100 trials), yielding an average of 2000 ± 317 trials (mean ± std) for each participant. Subjects received monetary compensation that depended on their performance (20€ + 0.01€ x N_corr_, where N_corr_ is the number of correct responses). The study was approved by the Ethics Committee for Clinical Research of the Barcelona Clinic Hospital.

### Analysis of human experimental data

We discarded 4 subjects with low performance (under 70%), and another 2 subjects whose median movement time was larger than 400ms (median across all trials from all subjects equal to 181 ms) (Chapman et al. 2010; Haith, Huberdeau, and Krakauer 2015). Fixation breaks and late responses were excluded from the analyses. We grouped all the trials from the remaining subjects together (except for the analysis of splitting time which was also performed separately for each subject). As for the rats, we analyzed only after-correct trials, for a total of 23,881 included trials. We analyzed the trajectories of the finger press from the starting point to the response point. Trajectories were obtained at a sample rate of 60 Hz, which were linearly interpolated to a 1000 Hz rate.

## Acknowledgements

The work was supported by the Spanish State Research Agency (RYC-2017-23231 to AH and AGD, PID2019-111629GB-I00 to AH, MMM and AGD; Severo Ochoa and María de Maeztu Program for Centers and Units of Excellence in R&D CEX2020-001084-M), Spanish Ministry of Economy and Competitiveness together with the European Regional Development Fund grant SAF2015-70324-R (to JR), European Research Council grant ERC-2015-CoG-683209 under the European Union’s Horizon 2020 (to JR); CERCA Programme/Generalitat de Catalunya. Part of the work was developed in the Centre Esther Koplowitz (Barcelona). J.R. appreciates the hospitality of the Grossman Center for Quantitative Biology and Human Behavior at the University of Chicago. MMM was supported by the National Institutes of Health under award number *1R01MH132172-01*.

## Supplementary figures

**Supplementary figure 1.**
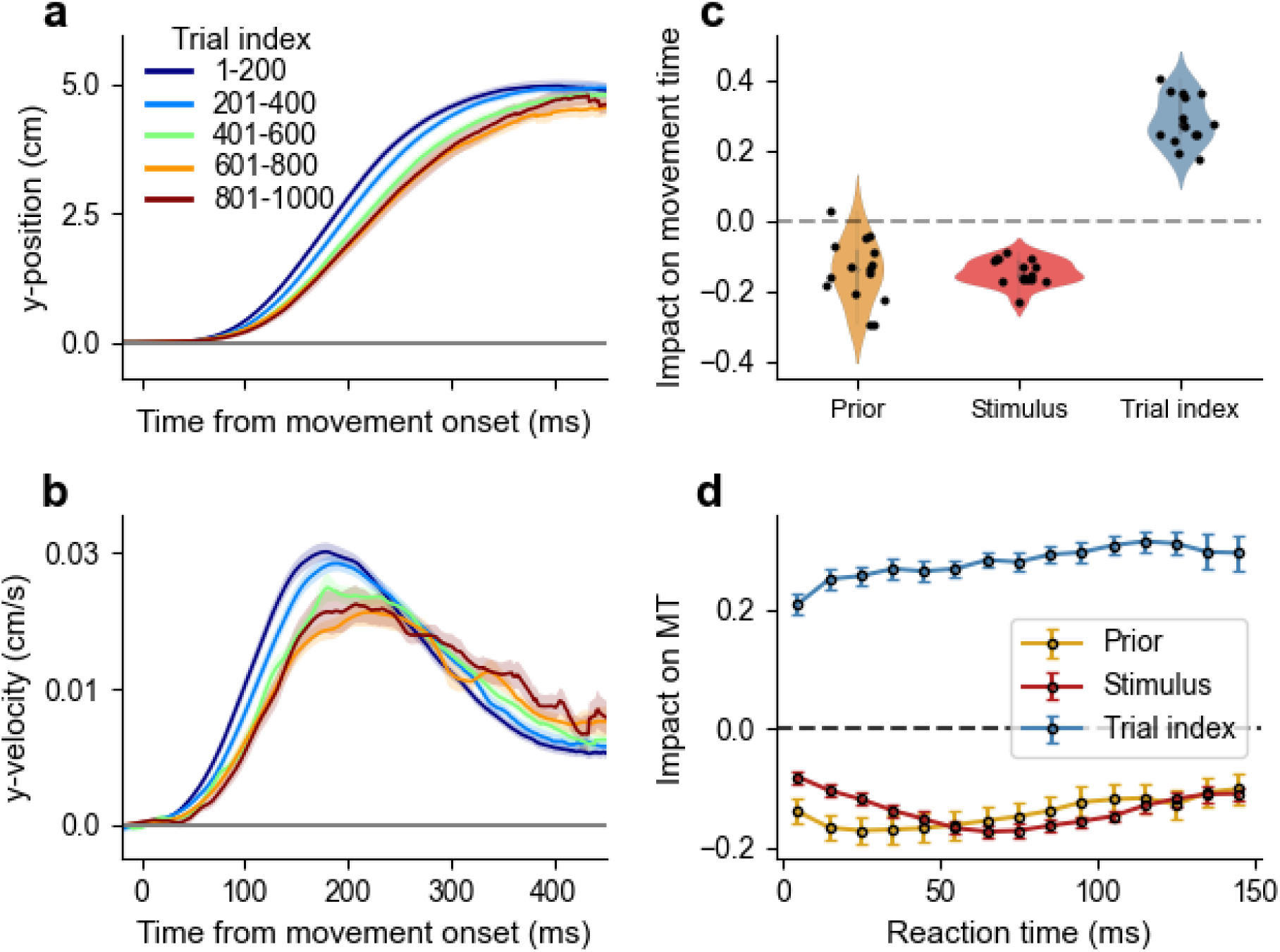
Trial index plays an important role for vigor. **a, b)** Average position in centimeters (a) and velocity in centimeters/s (b) in express responses with low prior (|z| < 25% percentile), conditioned on the trial index, grouped by bins of 200 trials. **c)** Regression weights of the movement time against prior evidence towards the response, stimulus evidence towards the response, and trial index. Points represent individual animals. **d)** Regression weights (mean ± s.e.m. across subjects) of the movement time against the trial index and the prior and stimulus evidence towards the response, for different bins of reaction time. The impact of stimulus evidence is lower for very short RTs (i.e. very short stimuli) while the impact of the prior decreases with increasing reaction time. This pattern is expected for an accumulation model where the movement time depends on the read-out of a decision variable whose offset and drift are determined by the prior and stimulus evidence, respectively.

**Supplementary figure 2.**
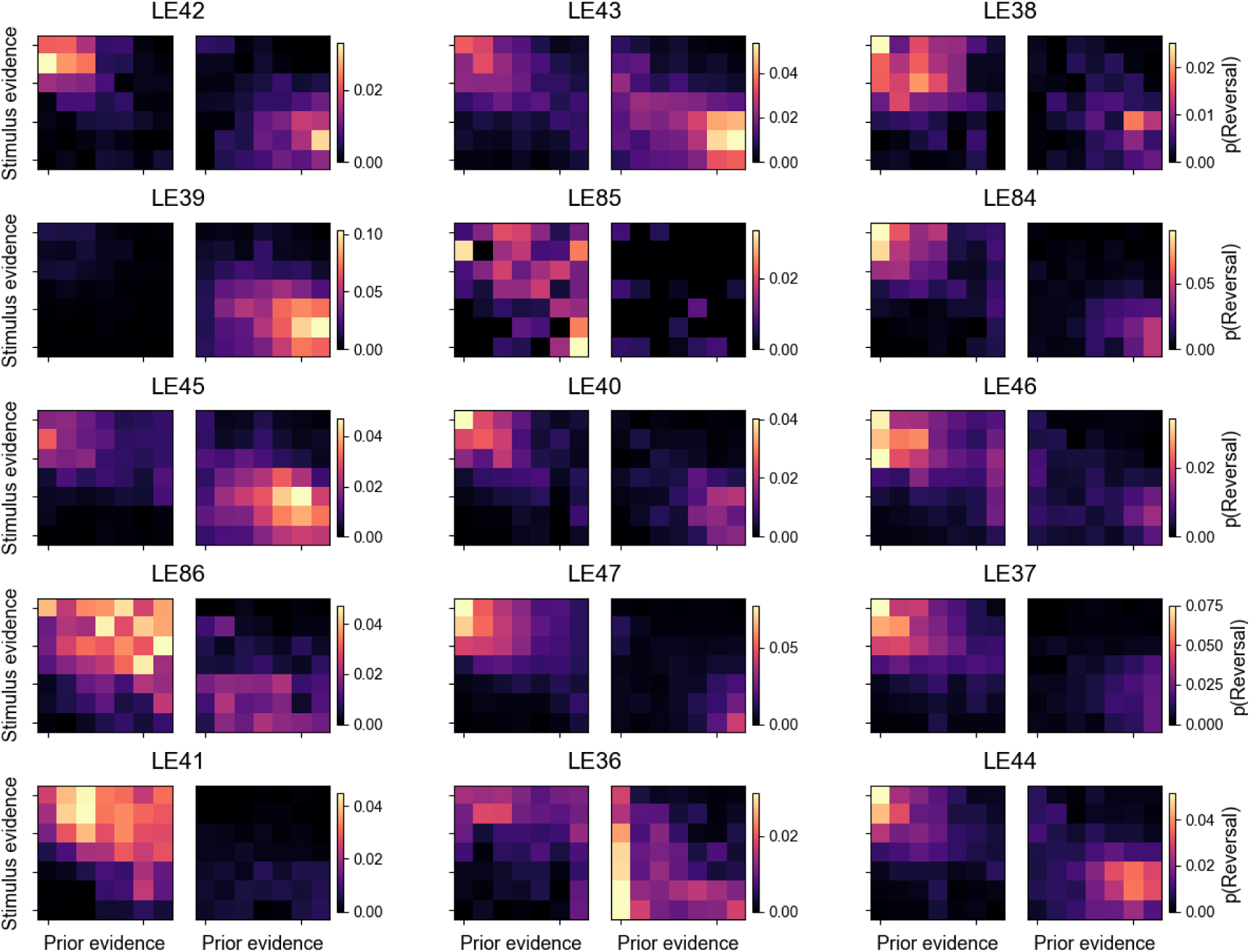
Proportion of left-to-right reversals (left matrices) and right-to-left reversals (right matrices) as a function of stimulus and prior evidence, for all animals.

**Supplementary figure 3.**
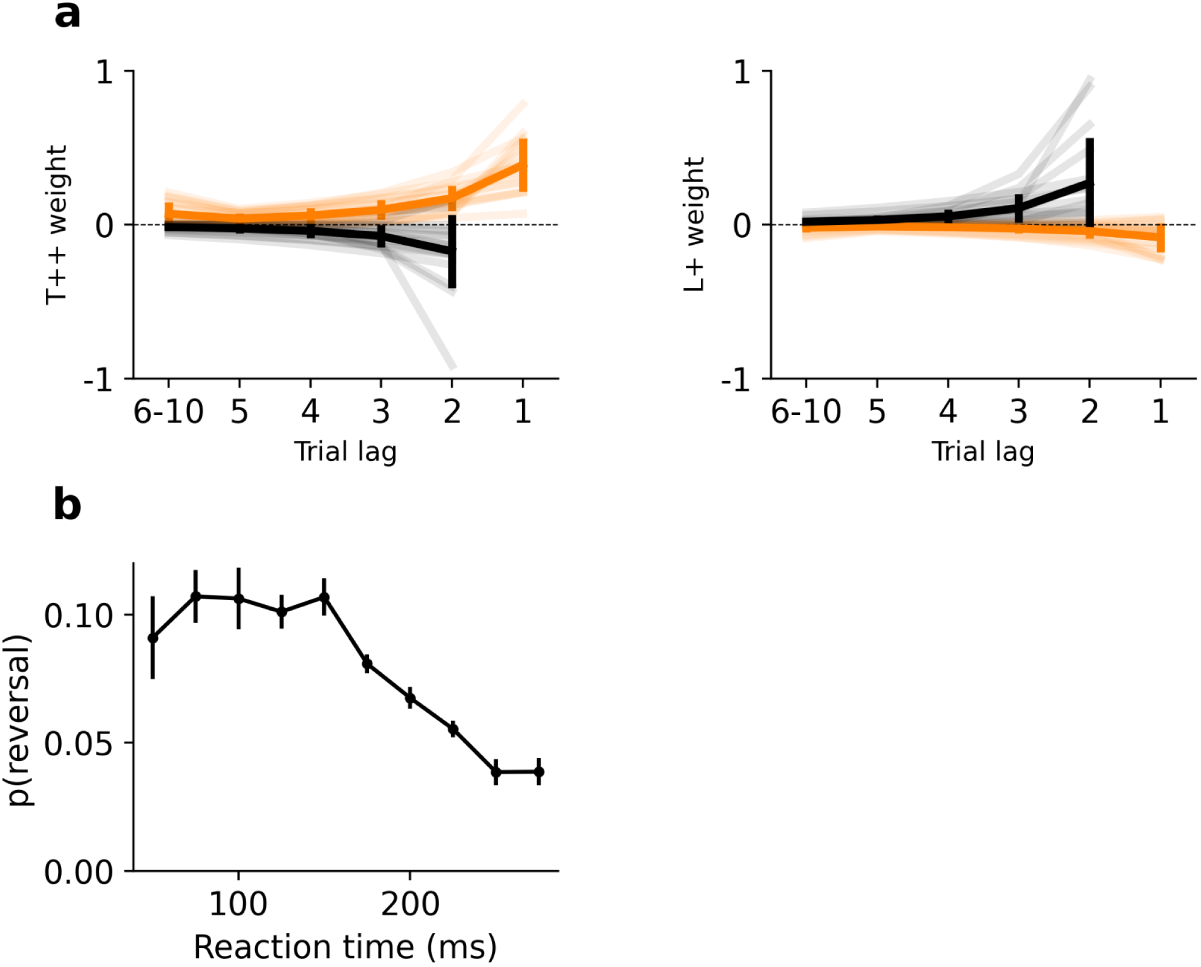
a) GLM transition (left) and lateral (right) weights for human participants. Orange: after correct trials; black: after error trials <u>(Hermoso-Mendizabal et al. 2020)</u>. Thin lines denote individual participants; black lines represent the average across participants (error bars: s.e.m.). **b)** Proportion of trajectory reversals as a function of reaction time for humans.

**Supplementary figure 4.**
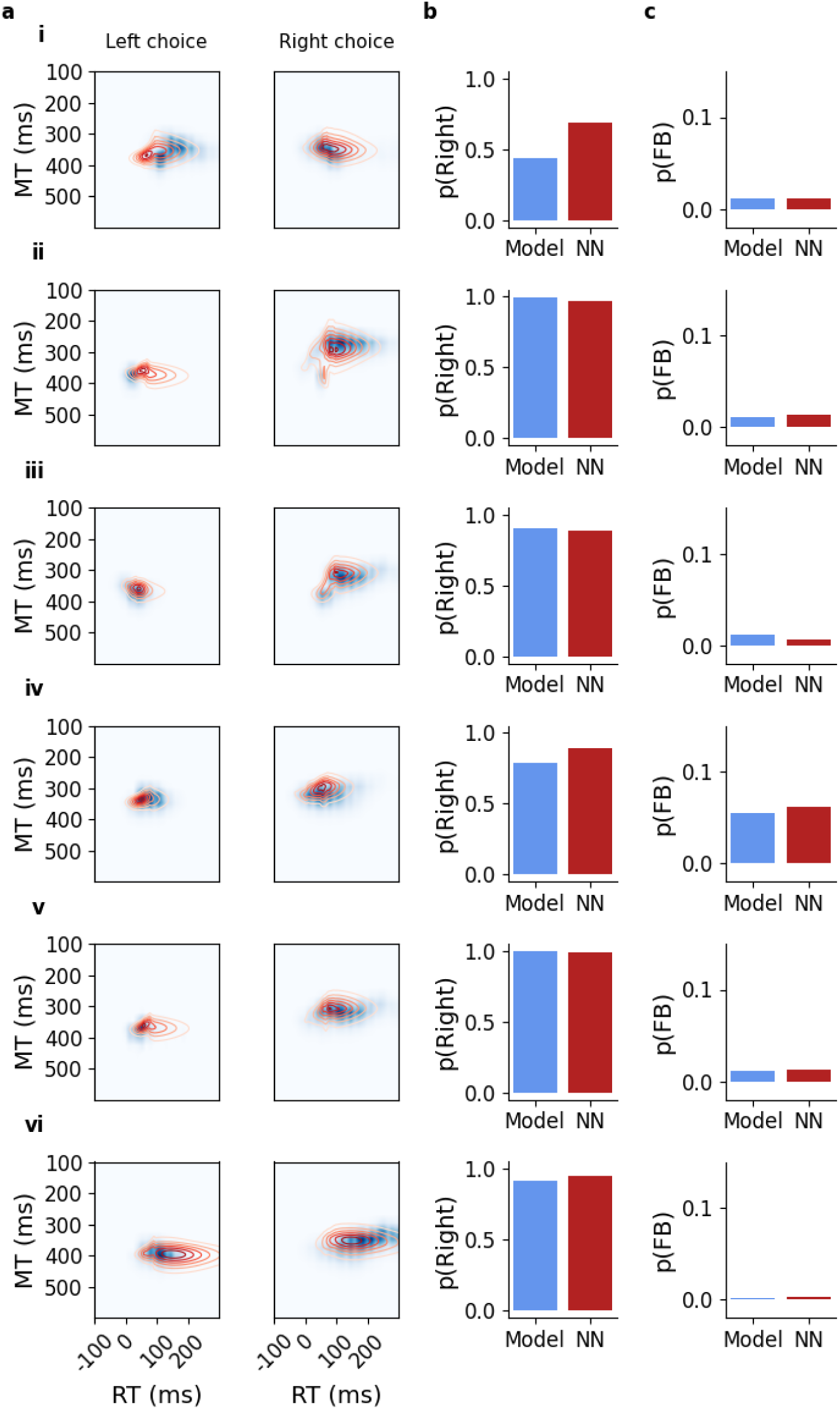
MNLE approximates the likelihood of the model. Each row represents fixed trial conditions (stimulus, prior and trial index). **a)** Likelihood for the model (blue-shaded matrix) based on 0.5M simulations and approximated likelihood obtained from the MNLE trained with 10M simulations (red contours); for left and right responses, respectively. **b-c)** Probability of rightward choice (b) and fixation break (c) for both model and MNLE. We used 6 prototypical experimental conditions: **i)** No stimulus evidence (s=0), prior towards the right (z=1.5), medium trial index (t.i.=400). **ii)** Low prior (z=0.05) and high stimulus strength (s=1) both towards the right choice, medium trial index (t.i.=400). **iii)** Prior (z=-1.5) towards the left choice, high stimulus (s=0.5) towards the right choice, medium trial index (t.i.=400). **iv)** Medium prior (z=0.5) and low stimulus (s=0.25), both towards the right choice, low trial index (t.i.=10). **v)** Strong prior (z=0.5) and strong stimulus (s=0.5), both towards the right choice, medium trial index (t.i.=400). **vi)** Medium prior (z=0.5) and low stimulus (s=0.25), high trial index (t.i.=800).

**Supplementary figure 5.**
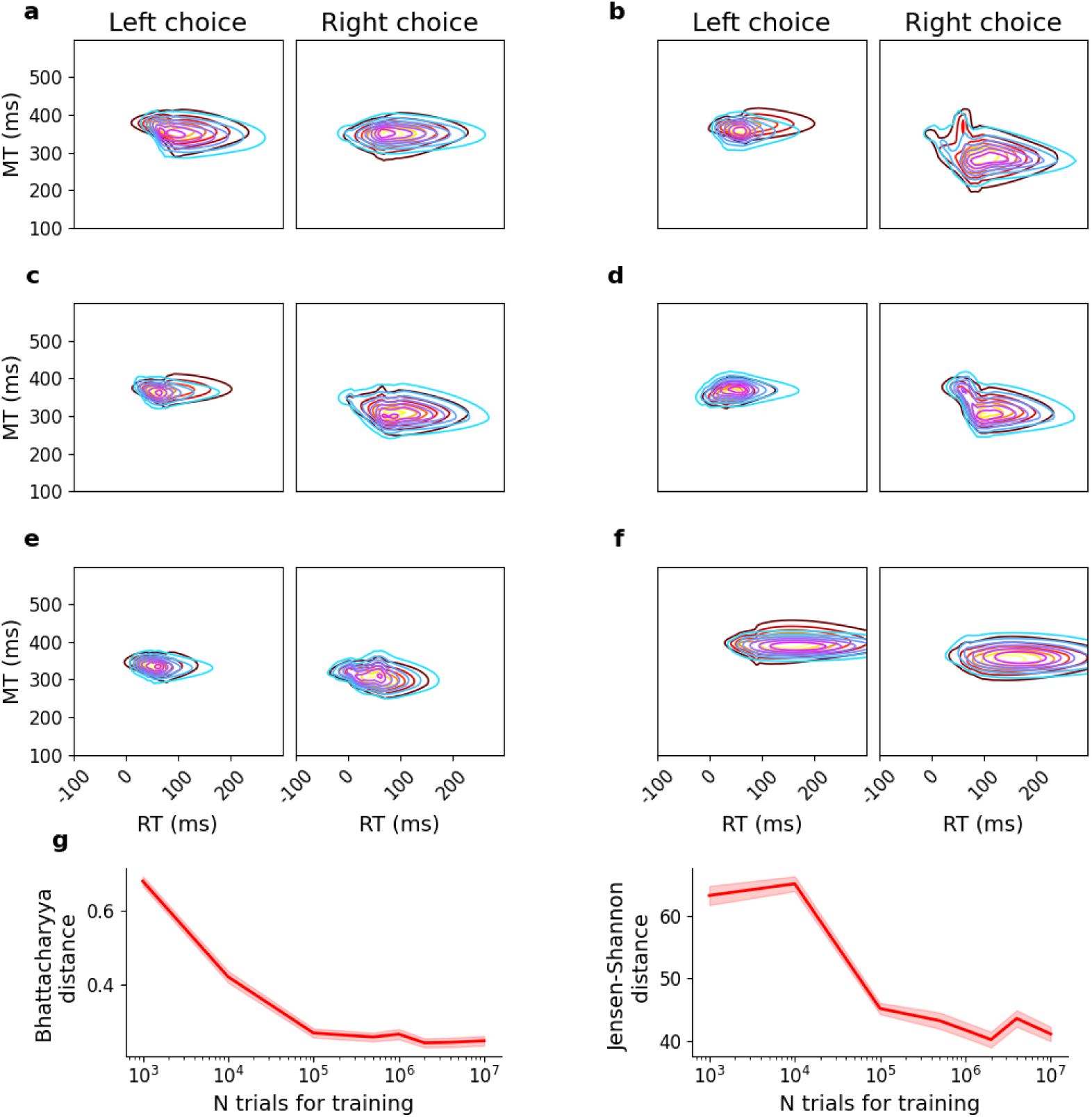
a-f) MNLE likelihood from two networks trained with a different set of 10M simulations. Same conditions as in the previous figure (Supplementary Fig. 4). **g)** Distance between the probability distributions from the network and the model, for different sizes of training sets for the neural network. We computed two distance metrics using 200.000 model simulations for 100 random trials. The metrics used were the Bhattacharyya distance (left panel), defined as 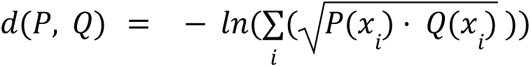; and the Jensen-Shannon distance (right panel), defined as 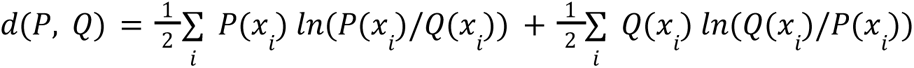; being *P* and *Q* the true and approximated likelihoods, respectively.

**Supplementary figure 6.**
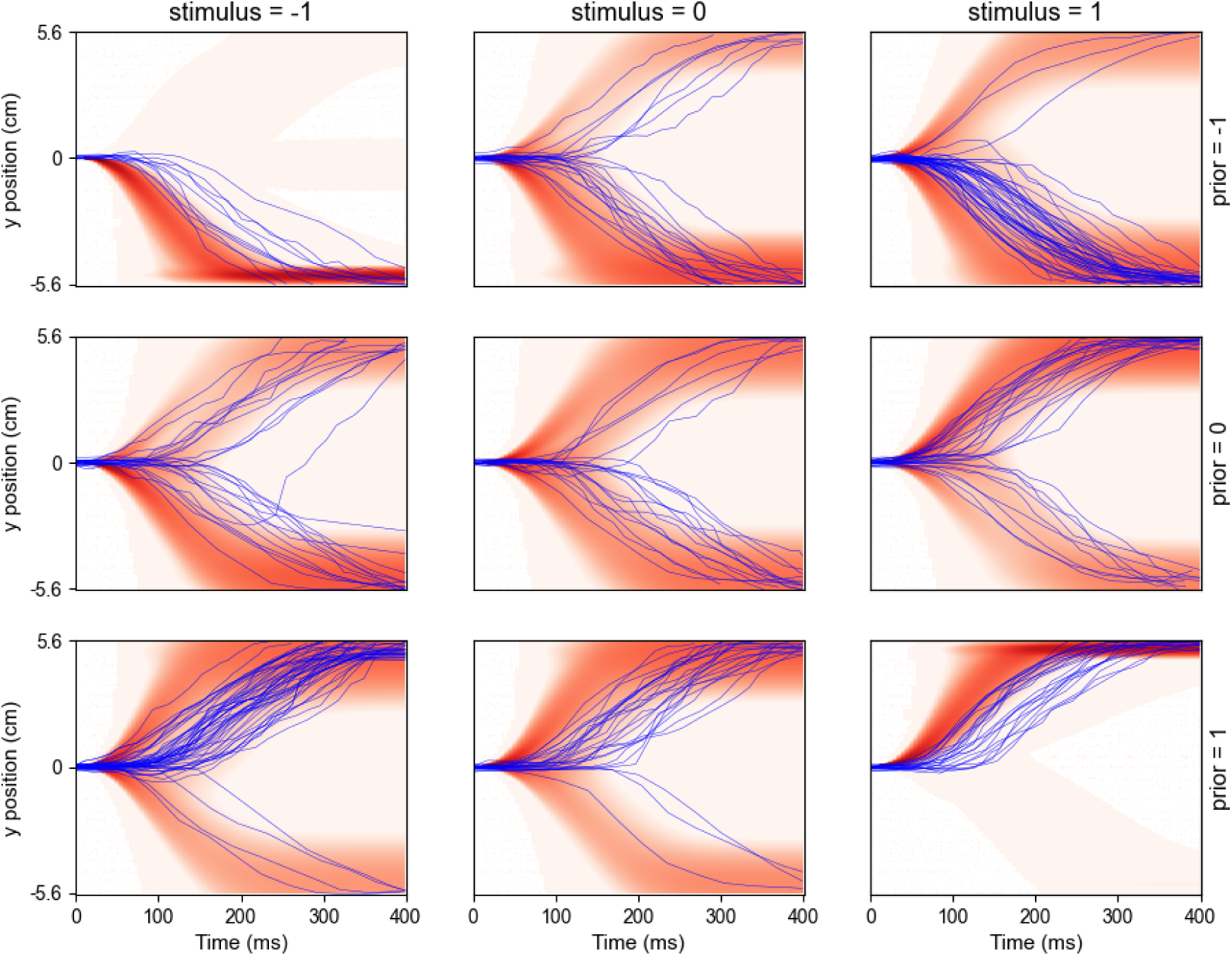
Rat trajectories for a random subset of 150 trials of animal LE43 (blue curves) plotted over the corresponding model trajectory density (red color plot), for different values of stimulus evidence (left column: stimulus=-1; middle column: stimulus=0; right column: stimulus=1) and prior evidence (top row: prior=-1, middle row: prior=0; bottom row: prior=+1). Model densities were obtained by drawing trajectories for 150000 simulated trials with model parameters estimated for the corresponding animal.

**Supplementary figure 7.**
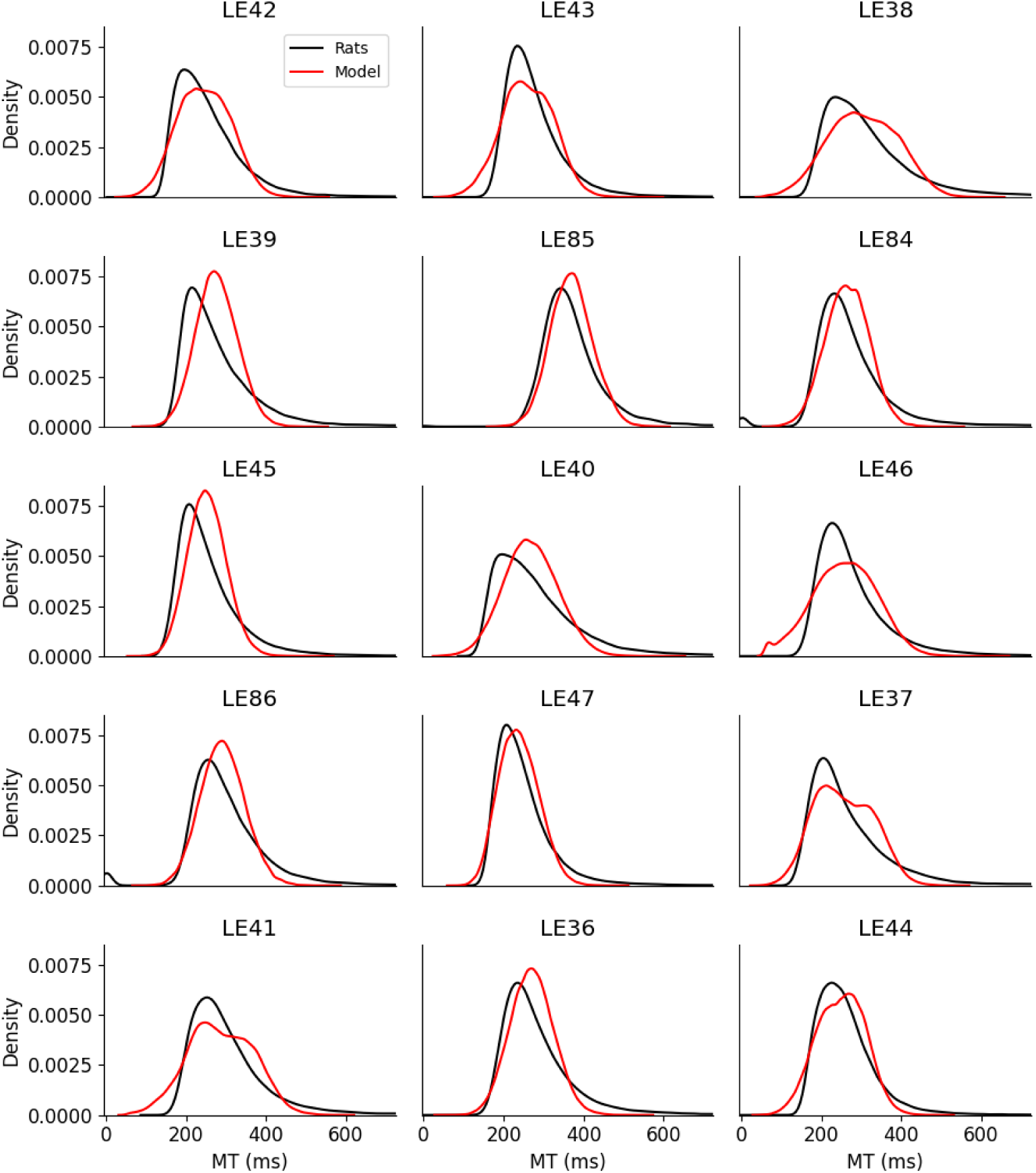
Rats and model MT distributions of each rat.

**Supplementary figure 8.**
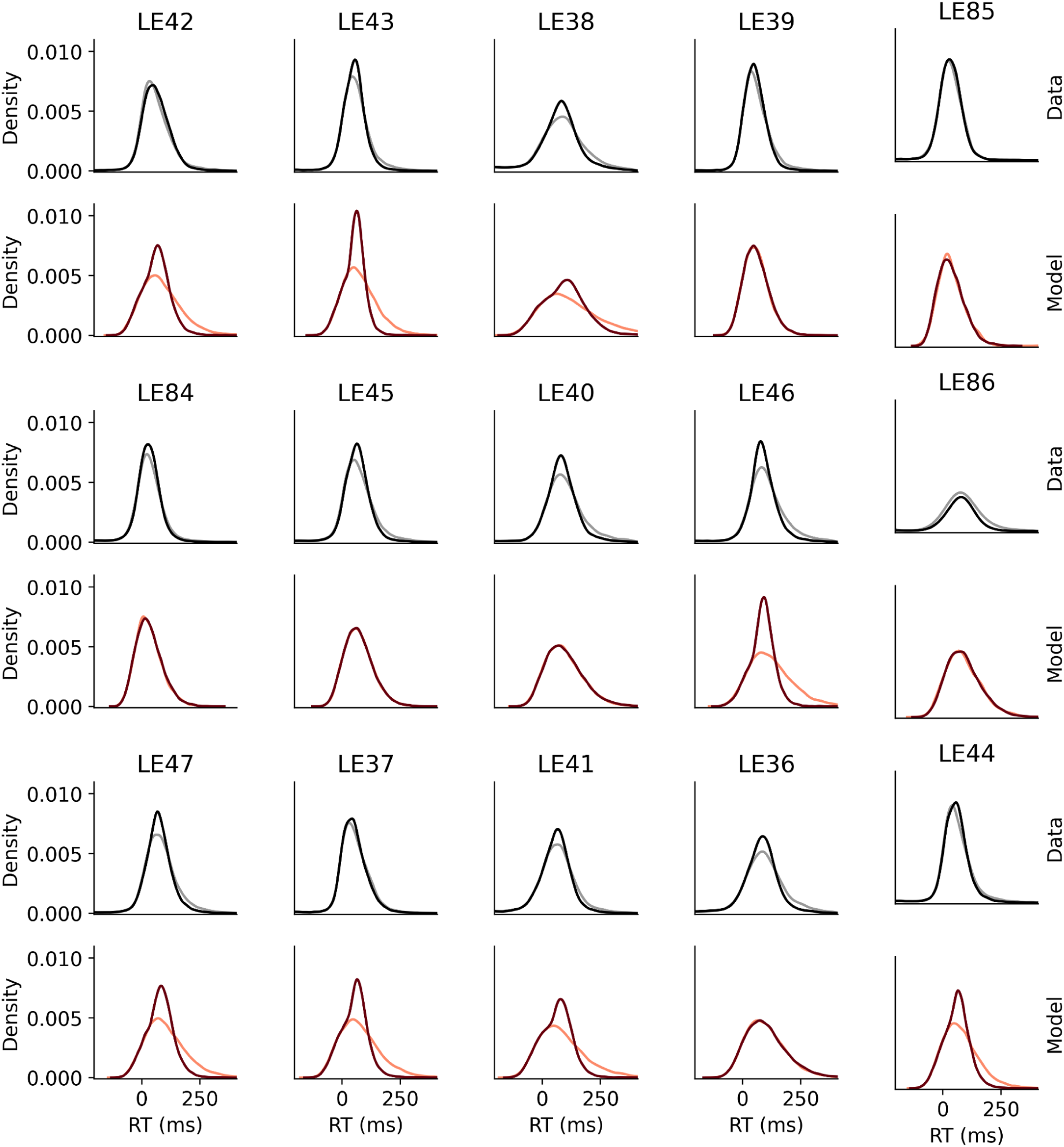
RT distributions from rats (gray/black lines) and model simulations (orange/brown lines), for different levels of stimulus strength (light: stimulus strength = 0; dark: stimulus strength = 1).

**Supplementary figure 9.**
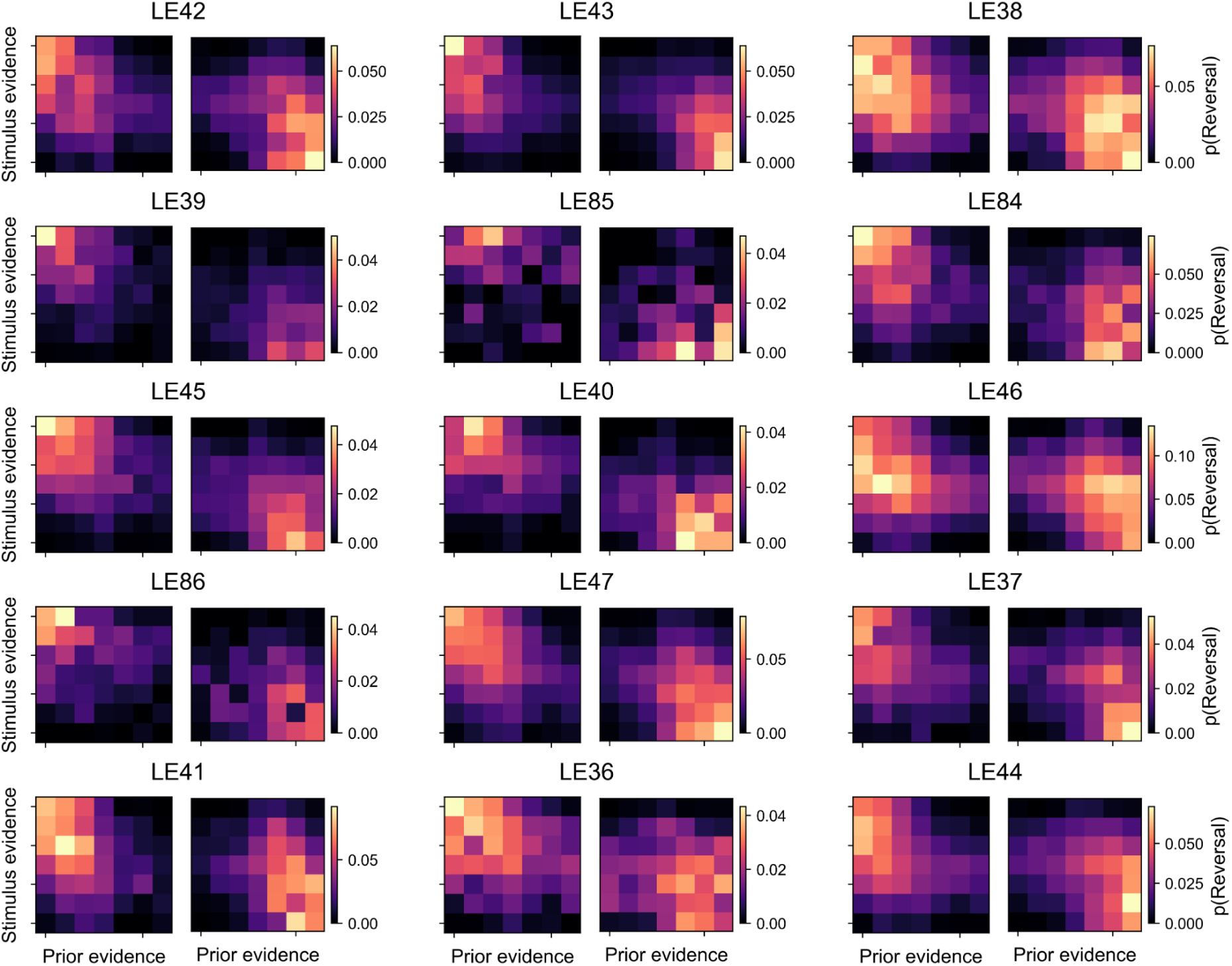
Proportion of left-to-right reversals (left panels) and right-to-left reversals (right panels) as a function of stimulus and prior evidence, for the model fitted to each individual rat.

**Supplementary figure 10.**
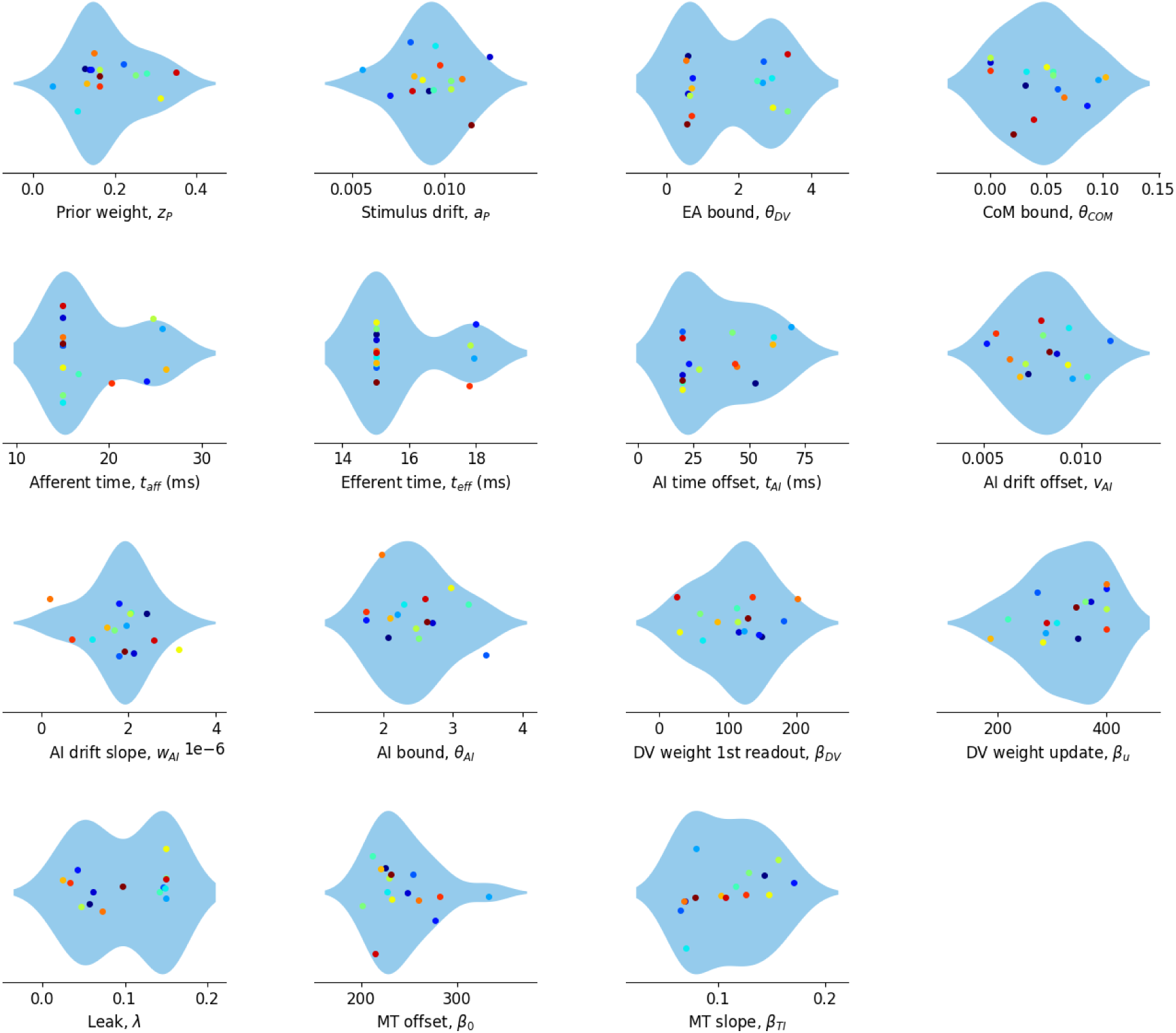
Fitted parameters for all rats. The parameter of the variance of the MT, σ_*MT*_, is not plotted, as it reached the upper bound for all rats (σ_*MT*_ = 20).

**Supplementary figure 11.**
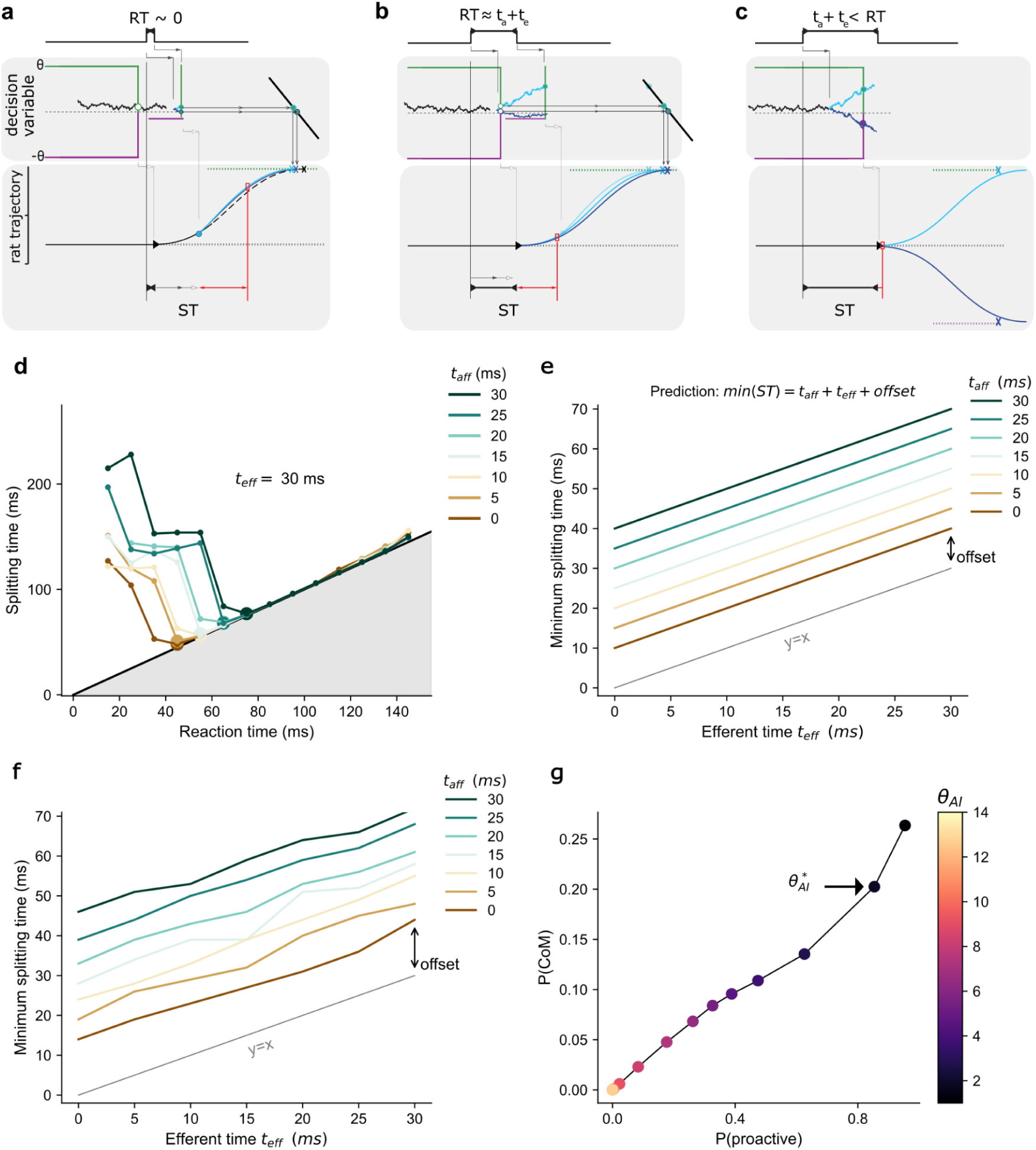
Model analysis of splitting times and changes of mind. **a-c)** Model schematics illustrating the dependence of splitting time (ST) on RT (Figure 6g). All panels show the evolution of the decision variable *x*(t) and the trajectory for two example stimuli (cyan and blue traces). ST can be defined as the interval between stimulus onset (black vertical line) and the moment when the two trajectories are statistically different (red vertical line, Methods). For RT < *t_aff_* + *t_eff_*, because of the afferent and efferent latencies, the stimulus can only be detected in the second read-out (a). After the updating of vigor, the subsequent trajectory depends on stimulus evidence (compare cyan and blue trajectories). In this condition, ST can be broken down into: ST ∼ RT + *t_aff_*+ *t_eff_* + *offset*, where the *offset* is the time it takes for the two updated trajectories to be distinguished (red double-arrow line), an interval that ultimately depends on the read-out difference between the two stimuli. For RT just above the value *t_aff_* + *t_eff_* (b) the stimulus starts to be detected in the first read-out making the ST ∼ RT + *offset*. As RT grows more, the first read-out can be more different between the two stimuli, generating very distinct trajectories that yield ST ≈ RT. **d)** ST as a function of reaction time for simulated model (averaged across animals), for various values of afferent time *t_aff_* (inset) and fixed efferent time *t_eff_* = 30 ms. The other parameters were set to the values obtained in fitting the model to one animal data. Larger dots indicate minimum splitting time for each value of *t_aff_*. Note that the minimum increases with *t_aff_*. **e)** Prediction of minimum splitting time, given by min(ST) = *t_aff_* + *t_eff_* + *offset*, as a function of efferent time. The offset was manually set to 10 ms. Gray line represents *y* = *x*. **f)** Minimum splitting time vs *t_eff_* obtained from model simulations shows good agreement with predictions in panel b. In simulations, the offset is caused by the noise in the evidence accumulation and motor processes and depends on the number of trials. **g)** Probability of CoM against probability of having a proactive trial, obtained by progressively slowing the model’s urgency signal (i.e. by increasing the boundary of the action initiation process), for the model fitted to the data of rat LE42. Note that CoMs vanish when there are no proactive trials (for very high values of the action initiation boundary). The value of the boundary found by maximum likelihood fitting yields the highlighted point 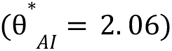.

**Supplementary figure 12.**
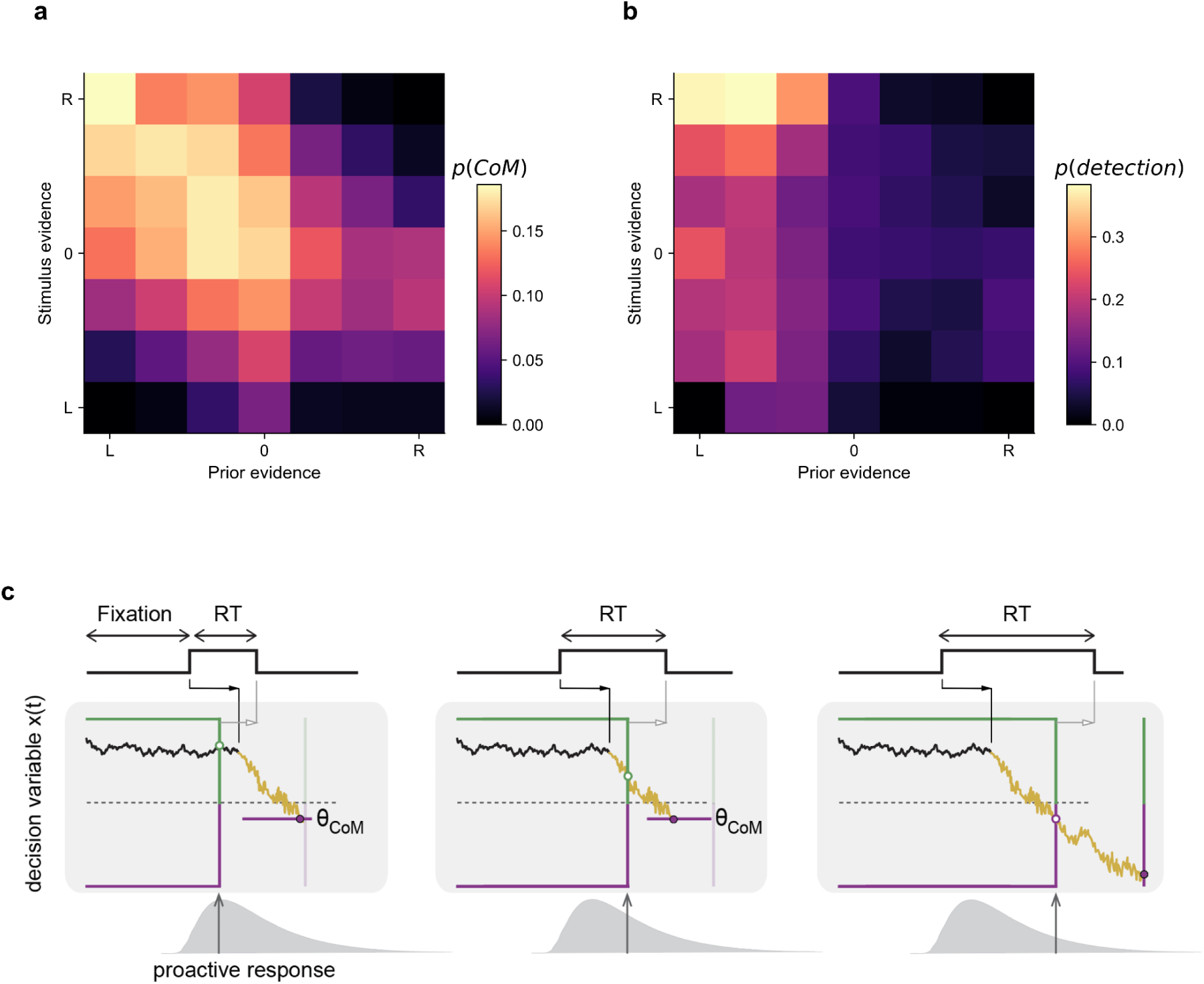
a-b) Proportion of left-to-right CoMs (**a**) and of CoMs that are detected as trajectory reversals (**b**) as a function of stimulus and prior evidence, for the model fitted to rat LE42. Note that the probability of left-to-right reversals is the product of these two quantities, since *p(reversal)=p(CoM)p(reversal|CoM).* CoMs are frequent when prior and stimulus evidence are incongruent, almost irrespective of the strength of evidence (or if anything, more frequently for weak/moderate evidence). However, CoMs are more often detected when evidence is strong. This is because the trajectory is more likely to cross the reversals detection threshold when the prior is strong and provides sufficient initial vigor. In turn, only a strong stimulus is capable of reversing a strong prior. **c)** Schematic of CoM generation for increasing reaction times (RT). When RT is small (express responses), the first read-out is determined by the prior (here, to the left) while the second read-out integrates contradictory evidence from the stimulus, leading to a CoM. By contrast, when the RT is large, the same evidence accumulation process leads to a first read-out which is already aligned with the stimulus (i.e. no CoM).

## Supplementary videos

**Supplementary Video 1.** Example responses from 8 rats showing two consecutive trials each: a correct non-CoM followed by a CoM (see label in the bottom right of the video). Every pair of consecutive trials is played first at real time and then replayed at a slowed velocity (x 0.25). Blue circles show the seven points tracked in the rat’s body used to estimate the general pose. The point at the base of the snout used to generate the orienting trajectories is shown in yellow followed by a “trail” composed of the location in the last 10 frames. The ninth tracked point at the tip of the snout is not shown for clarity. At the bottom right of the video, during the CoM trial, “y_rev” indicates the reversal point of the trajectory in pixels (detection threshold was set at 8 pixels (i.e. 0.56 cm).

